# Female-specific m6A remodeling in the liver correlates with post-transcriptional metabolic adaptation to high fat diet

**DOI:** 10.64898/2026.06.19.733425

**Authors:** Sofia V. Krylova, Maxwell Horton, Giorgia Bucciarelli, Li Liu, Jacob Berrigan, Ronald Cutler, Kartik Chandran, Nathaniel W. Snyder, Toma Tebaldi, Simone Sidoli, Kamini Singh, Jeffrey E. Pessin

## Abstract

Sex differences strongly influence susceptibility to metabolic dysfunction-associated steatotic liver disease (MASLD), yet the regulatory mechanisms underlying these differences remain incompletely understood. To examine sex-specific hepatic adaptation to a high-fat (HF) diet mouse model of MASLD, we integrated proteomics, transcriptomics, and Oxford Nanopore direct RNA sequencing for transcriptome-wide m6A profiling in male and female mouse livers. Female mice were relatively protected from HF diet–induced hepatic steatosis and exhibited distinct proteome remodeling enriched for peroxisomal pathways. In contrast, transcriptomic responses in females were dominated by inflammatory signatures and did not recapitulate the metabolic adaptations observed at the protein level, revealing extensive RNA–protein discordance and post-transcriptional remodeling. Integrated RNA–protein analyses identified female-specific amplification of peroxisomal proteins despite modest transcript-level changes. HF diet also induced sex-specific remodeling of m6A RNA methylation and altered regulation of the m6A methylation system. Notably, reduced 3′ UTR m6A methylation of peroxisomal transcripts inversely correlated with increased protein abundance relative to RNA expression in female mice. Together, these findings implicate m6A-associated post-transcriptional regulation in sex-specific hepatic adaptation to HF diet exposure and the basis for discordance between many of the mRNAs and proteins in the liver.

## Introduction

Metabolic dysfunction-associated steatotic liver disease (MASLD) and its progressive forms are strongly associated with obesity, insulin resistance, and chronic nutrient excess ^1,2^. Marked sex differences exist in MASLD susceptibility and progression, with premenopausal females generally exhibiting relative protection from hepatic steatosis compared to males ^3–5^. These differences suggest the presence of sex-specific mechanisms that regulate hepatic lipid metabolism and adaptation to nutritional stress. Although sexually dimorphic transcriptional programs governing hepatic lipid metabolism have been extensively studied as mechanisms of differential MASLD susceptibility, including regulation by the GH/STAT5 and BCL6 axes, transcript abundance alone is often insufficient to predict protein abundance or metabolic phenotypes in the liver ^6,7^. Indeed, extensive RNA–protein discordance has been reported across multiple metabolic pathways, highlighting the importance of regulatory mechanisms beyond transcription ^8–10^. Increasing evidence indicates that post-transcriptional mechanisms, including RNA modifications, RNA stability, and translational control, play major roles in shaping hepatic metabolic responses ^11–13^.

Among these post-transcriptional regulatory pathways, N6-methyladenosine (m6A) is among the most abundant internal mRNA modifications in eukaryotic transcripts ^14,15^. m6A regulates multiple aspects of RNA metabolism, including mRNA stability, translation, localization, and degradation, through interactions with m6A reader proteins (15). m6A deposition is catalyzed by methyltransferase complexes containing METTL3 and METTL14, whereas demethylases such as FTO and ALKBH5 remove these marks ^16^. Altered m6A regulation has been implicated in obesity, hepatic steatosis, lipid metabolism, as well as sex-dimorphic metabolic traits, including through effects on transcript turnover and hepatic lipid regulation ^11,17,18^. However, whether diet-induced m6A remodeling contributes to sex-dimorphic hepatic proteome remodeling and differential susceptibility to steatosis remains poorly understood, particularly since the relationship between m6A regulation and protein-RNA discordance in male and female liver has not been systematically investigated.

Peroxisomes contribute to hepatic lipid homeostasis and MASLD susceptibility through roles in very long chain and branched chain fatty acid oxidation, ether lipid synthesis, bile acid precursor processing, and redox metabolism ^19–21^. However, peroxisomal pathways appear to have context-dependent roles in MASLD, with evidence implicating both adaptive lipid-handling functions and disease-associated peroxisomal dysfunction ^22–24^. Despite the established importance of peroxisomes in hepatic metabolism, whether sex-specific regulation of peroxisomal pathways contributes to female protection from diet-induced steatosis remains unclear. In particular, whether epi-transcriptomic mechanisms participate in sex-specific post-transcriptional regulation of peroxisomal proteins during metabolic stress is unknown. In this study, we integrated transcriptomics, quantitative proteomics and Oxford Nanopore direct RNA sequencing to define sex-specific hepatic responses to HF diet in mice.

## Results

### Female mice are relatively protected from high–fat diet–induced hepatic steatosis

Sex differences in susceptibility to MASLD are well established, with adult female humans and mice frequently showing reduced hepatic steatosis in response to obesogenic diets compared to males ^3,25–27^. To investigate the molecular mechanisms underlying this protective effect, we first validated sexually dimorphic responses in our experimental conditions in a mouse model of MASLD induced by high-fat diet feeding.

Female (n=4) and male (n=4) C57BL/6J mice were maintained on either normal chow (NC) or high-fat (HF) diet from 6 to 16 weeks of age (Figure 1A). Both sexes exhibited increased weight gain on HF diet relative to NC controls (Figure 1B). Analysis of body weight at the experimental endpoint (16 weeks of age; 10 weeks on diet) by two-way ANOVA revealed a significant sex × diet interaction (F statistic = 8.27, degrees of freedom = 1.12; P = 0.0139), indicating that the effect of HF diet on body weight differed between males and females. Post hoc Fisher’s comparisons showed significant increases in body weight under HF diet in both males and females (males NC vs HF: mean difference = −17.73 g, P < 0.0001; females NC vs HF: mean difference = −11.60 g, P < 0.0001), with males exhibiting greater body weight than females under both NC and HF diet conditions (NC: mean difference = 7.125 g, P = 0.0005; HF: mean difference = 13.25 g, P < 0.0001).

**Figure 1:**
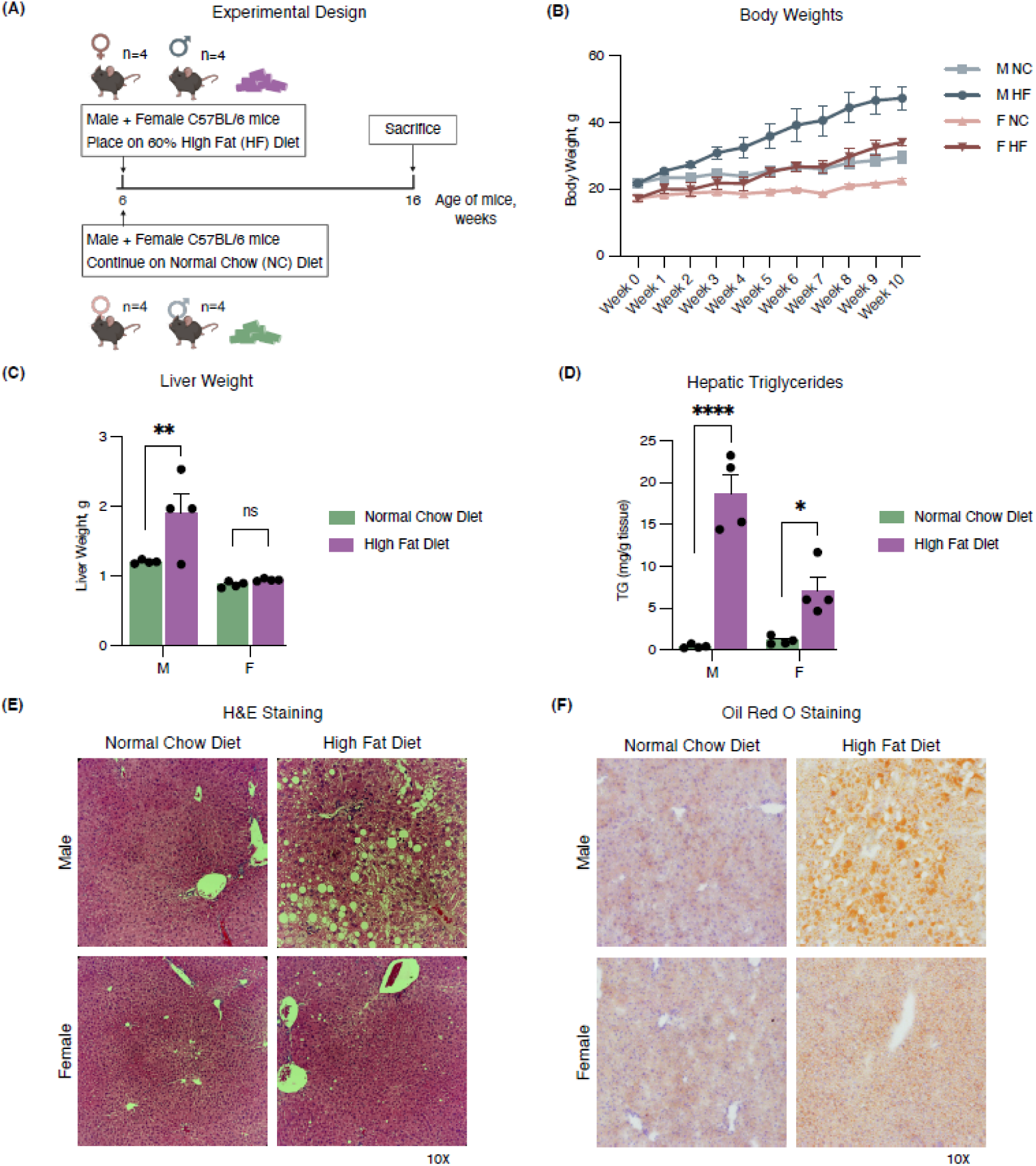
Female mice are relatively protected from high-fat diet–induced hepatic steatosis. (A) Schematic of experimental design. Male (M) and female (F) C57BL/6 mice (n = 4 mice per group) were placed on high-fat (HF) diet or maintained on normal chow (NC) from 6 to 16 weeks of age before whole liver collection. (B) Body weight progression over time in M and F mice on NC versus HF diets. (C) Liver weights at collection in M and F mice on NC versus HF diets. (D) Hepatic triglyceride (TG) content in M and F mice on NC versus HF diets. (E) Representative haematoxylin and eosin (H&E) staining of liver sections (10× magnification) in M and F mice on NC versus HF diets. (F) Representative Oil Red O staining of lipid accumulation (10× magnification) in M and F mice on NC versus HF diets. Data are presented as mean ± s.e.m.; n = 4 mice per group. Statistical significance was assessed using two-way ANOVA followed by multiple comparisons testing (Šídák correction), comparing NC and HF conditions within each sex.

Although the HF diet increased total body weight in both sexes, hepatic lipid accumulation was more pronounced in males than in females. Consistent with this, liver weights were increased in HF-fed mice in a sex-dependent manner (Figure 1C). Two-way ANOVA revealed significant effects of sex and diet, as well as a significant sex × diet interaction (F(1,12) = 5.0, P = 0.0434). Post hoc analysis showed a significant increase in liver weight in HF-fed males compared to NC controls (Šídák-adjusted P = 0.0083), whereas the increase in females did not reach significance (P = 0.9342).

Similarly, although HF diet increased hepatic triglyceride (TG) content in both sexes, the magnitude of TG accumulation was significantly greater in males than in females (Figure 1D). Two-way ANOVA revealed significant effects of diet and sex, as well as a significant sex × diet interaction (F(1,12) = 20.0, P = 0.0008), with males exhibiting a substantially larger increase in TG content than females (mean difference NC vs HF: −18.25 vs −5.96; Šídák-adjusted P < 0.0001 and P = 0.0193, respectively). Histological analysis further supported these findings. H&E staining revealed pronounced macrovesicular steatosis in male livers following HF feeding, whereas female livers did not exhibit significant lipid droplet accumulation (Figure 1E). Oil Red O staining further confirmed reduced lipid accumulation in HF-fed females relative to males (Figure 1F). Together, these findings demonstrate that our mouse model recapitulates the sexually dimorphic susceptibility to high-fat diet–induced hepatic steatosis, with females exhibiting relative protection compared to males.

### Female mice on a high–fat diet display unique proteome remodeling and enrichment of the peroxisome pathway

To identify molecular pathways underlying the sex-specific differences in hepatic steatosis, we performed quantitative proteomic profiling of liver tissue from male and female mice under NC and HF diet conditions. Proteomics quality control and filtering metrics are summarized in Supplementary Figure S1A-D. Principal component analysis (PCA) revealed tight clustering of biological replicates within each condition, indicating high reproducibility of the dataset (Figure 2A). The primary axis of variation (PC1; 23.0% of variance explained) separated samples by sex, whereas the secondary axis (PC2; 17.8%) distinguished HF vs NC diets within each sex. These findings indicate that sex is the dominant source of proteomic variation regardless of the diet, whereas the secondary axis captures HF diet–induced proteomic remodeling within males and females. Accordingly, all subsequent analyses focused on within-sex HF vs NC comparisons to resolve diet-induced effects without confounding from the larger baseline proteomic differences between sexes.

**Figure 2:**
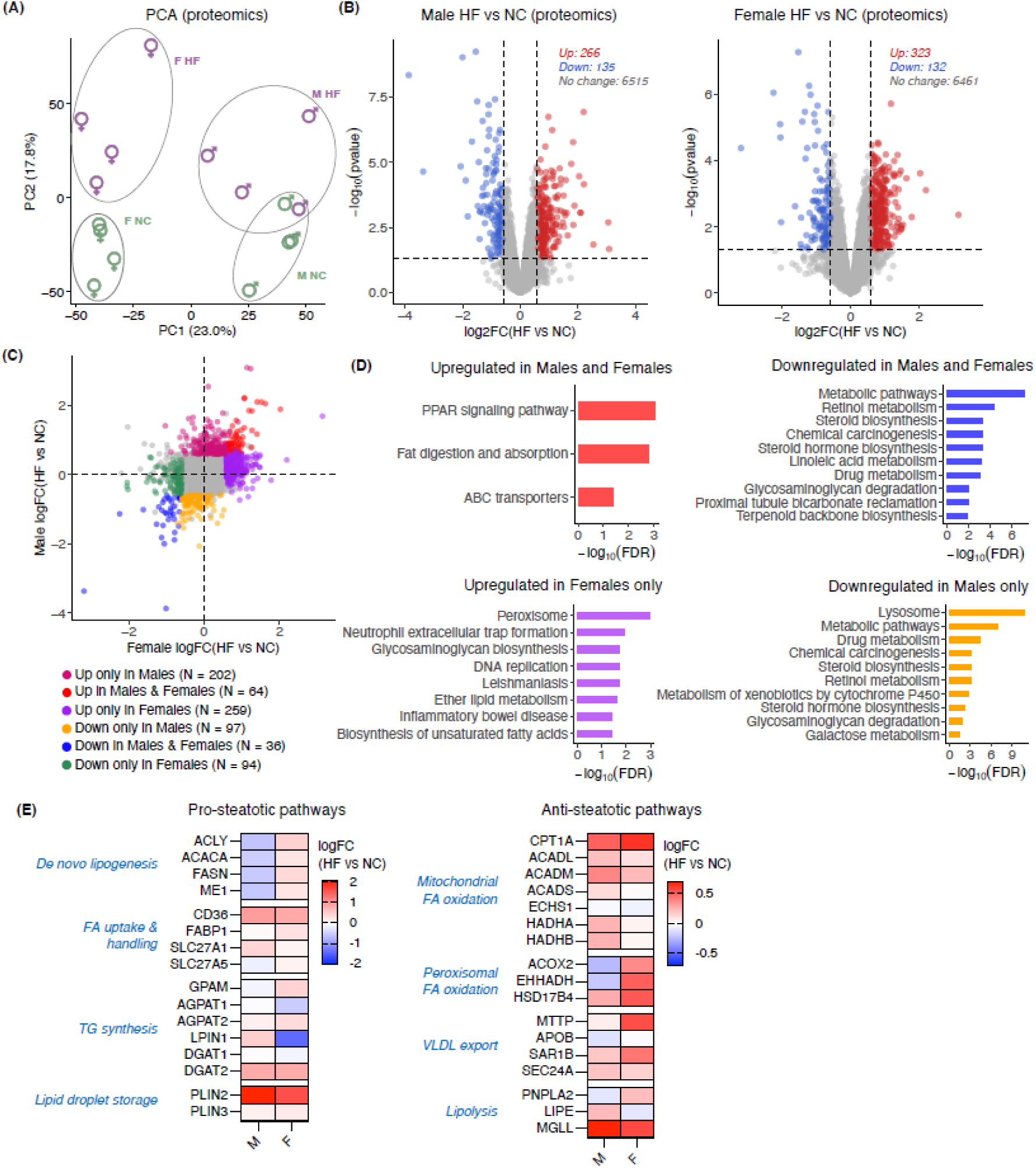
Female mice on HF diet display unique proteome remodeling and enrichment of the peroxisome pathway. (A) Principal component analysis (PCA) of liver proteomes from male (M) and female (F) mice fed normal chow (NC) or high fat (HF) diets (n = 4 mice per group). Each point represents a biological replicate. (B) Volcano plots showing differential protein abundance in M (left) and F (right) mice on HF vs NC diets. Differential expression was calculated using limma; proteins with |logFC| > 0.585 and p-value < 0.05 are highlighted. (C) Comparison of logFC (HF vs NC) between M and F, indicating proteins that are significantly up- and downregulated in males only, females only, or both. (D) KEGG pathway enrichment analysis (ShinyGO v0.77) of differentially regulated proteins in each category from (C). Downregulated in females only and upregulated in males only groups showed no significant enrichment. Top 10 with an enrichment threshold of FDR < 0.05 are displayed and ranked by –log (FDR). (E) Heatmaps of proteins involved in pro-steatotic and anti-steatotic pathways in the liver. Color scale represents limma logFC (HF vs NC) values for M and F.

Differential expression analysis using limma revealed substantial proteomic remodeling in response to HF diet in both males and females (Figure 2B). Across the male and female datasets, 6916 total proteins were identified. Using thresholds of |logFC| > 0.585 (1.5-fold change) and p-value < 0.05, we defined 266 upregulated and 135 downregulated proteins in males, and 323 upregulated and 132 downregulated proteins in females. While the volcano plots demonstrate that HF diet induces predominantly upregulated rather than downregulated proteomic changes in both sexes, they do not resolve what changes are shared versus those that are sex-specific.

To directly compare diet-induced responses between females and males, we plotted protein log2 fold changes (protein logFC) (HF vs NC) between sexes and categorized the significantly changing proteins (|logFC| > 0.585 and p-value < 0.05) into six groups: upregulated in both males and females, upregulated in males only, upregulated in females only, downregulated in both males and females, downregulated in males only, and downregulated in females only (Figure 2C). Interestingly, while both the upregulated and downregulated groups showed overlap between males and females (up in both: n = 64; down in both: n = 36), there were more than twice as many proteins uniquely upregulated or downregulated within each sex compared to proteins commonly regulated between sexes (up only in males: n = 259; up only in females: n = 202; down only in males: n = 97; down only in females: n = 94). These results demonstrate that livers of females versus males display largely unique adaptations to HF diet at the proteome level.

To determine the functional significance of these sex-specific proteomic changes, we performed KEGG pathway enrichment analysis on each category of proteins defined in figure 2C (Figure 2D). Proteins downregulated uniquely in females or upregulated uniquely in males did not show significant pathway enrichment and are therefore not displayed. Proteins upregulated in both males and females demonstrated enrichments for pathways related to lipid metabolism and transport, including PPAR signaling, fat digestion and absorption, and ABC transporters, consistent with a shared hepatic response to increased lipid delivery and processing under a high-fat diet. Pathways commonly downregulated in both sexes included broad metabolic and biosynthetic processes, such as steroid and retinol metabolism. Interestingly, the top significant enrichment for proteins upregulated uniquely in HF-fed females was the peroxisome pathway. Peroxisomes are known to play a significant role in hepatic fatty acid metabolism, including the α-oxidation of branched-chain fatty acids and the β-oxidation of very-long-chain fatty acids, as well as the synthesis bile acids and plasmalogen phospholipids ^28^.

It has been previously shown that impaired peroxisomal activity can exacerbate hepatic lipid accumulation and promote progression of steatotic liver disease ^20,25,26^. To assess the potential contribution of peroxisomal fatty acid oxidation pathways to female steatosis protection, we examined HF diet–induced changes in key proteins involved in major pro-steatotic (de novo lipogenesis, fatty acid uptake, triglyceride synthesis, lipid storage) and anti-steatotic (lipolysis, mitochondrial and peroxisomal fatty acid oxidation, and lipid export) processes in females and males (Figure 2E). Across most pathways, the direction and magnitude of protein abundance changes were broadly similar between sexes or occurred in directions inconsistent with female-selective steatosis protection. For example, several proteins involved in de novo lipogenesis were increased in females and decreased in males under HF diet conditions. Only two pathways displayed HF diet–induced protein changes consistent with reduced steatosis in females: peroxisomal fatty acid oxidation and VLDL export. Key proteins involved in peroxisomal fatty acid oxidation (ACOX2, EHHADH, HSD17B4) exhibited coordinated upregulation selectively in females, whereas proteins involved in VLDL export showed greater induction in females than in males. VLDL export has been previously demonstrated to be affected by estradiol and potentially implicated in protection against hepatic steatosis in females ^29–31^. In contrast, enhanced peroxisomal fatty acid oxidation has not previously been studied as a female-specific adaptive response to HF diet–induced steatosis.

These findings suggest that female protection against hepatic steatosis may involve previously unrecognized protein-level metabolic adaptations, particularly within peroxisomal pathways. Therefore, we next sought to determine how these sex-specific proteomic responses are regulated. Given the recently documented discordance between RNA and protein abundances in the liver ^8,10^, particularly among metabolic genes, we performed transcriptomic profiling to assess whether the observed proteomic remodeling could be explained at the transcriptional level.

### Transcriptome-level changes do not recapitulate sexually dimorphic proteomic patterns in response to high–fat diet

To determine whether the sex-specific proteomic remodeling observed under HF diet is reflected at the transcriptional level, we performed RNA-sequencing (RNA-seq) of liver tissue from male and female mice. RNA-seq quality control and filtering metrics are summarized in Supplementary Figure S2A-F. Principal component analysis (PCA) of liver transcriptomes revealed tight clustering of biological replicates within each condition, indicating high reproducibility of the dataset (Figure 3A). The primary axis of variation (PC1; 43.36% of variance explained) separated samples by sex, whereas the secondary axis (PC2; 16.35%) distinguished diet within each sex. Notably, while PC2 accounted for a similar proportion of variance in both the transcriptomic and proteomic datasets (∼16–18%), the contribution of PC1 was substantially greater in RNA-sequencing compared to proteomic data (43.36% vs 23.0%, respectively), suggesting stronger sex-associated separation in the transcriptomic compared to the proteomic dataset.

**Figure 3.**
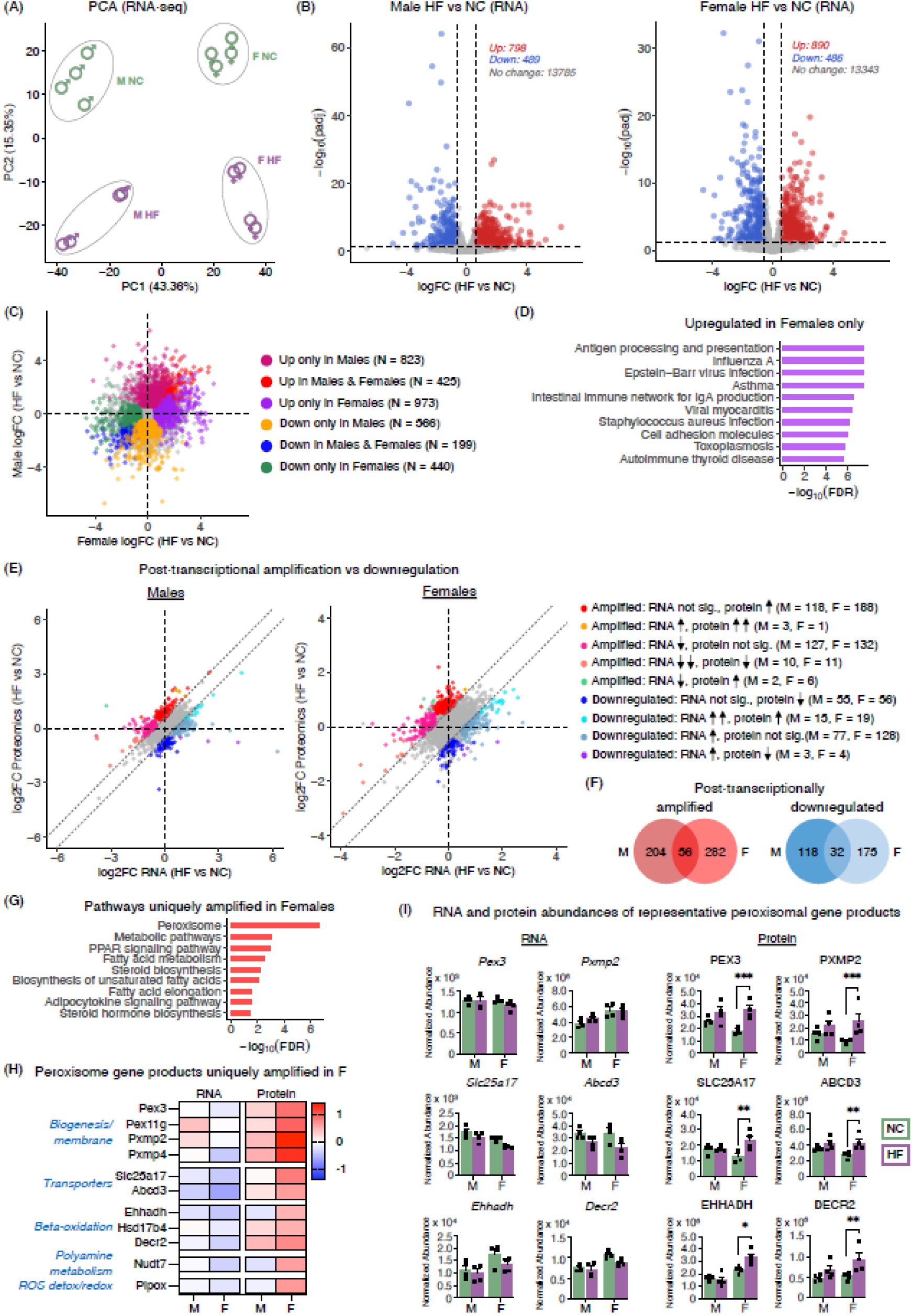
Transcriptome-level changes do not recapitulate sexually dimorphic proteomic patterns in response to HF diet. (A) Principal component analysis (PCA) of liver transcriptomes from male (M) and female (F) mice fed normal chow (NC) or high-fat (HF) diets (n = 4 mice per group). Each point represents a biological replicate. (B) Volcano plots showing differential gene expression in M and F mice on HF versus NC diets. Differential expression was calculated using DESeq2; genes with |logFC| > 1 and adjusted p-value (padj) < 0.05 are highlighted. (C) Comparison of RNA logFC values (HF vs NC) between M and F, indicating genes significantly upregulated or downregulated in males only, females only, or both, based on the same thresholds as in (B). (D) KEGG pathway enrichment analysis using ShinyGO v0.77 of differentially regulated genes in the Upregulated in Females Only category from (C). The top 10 pathways meeting an FDR < 0.05 threshold are shown and ranked by −log (FDR). (E) Comparison of RNA and protein logFC values (HF vs NC) highlighting gene products classified as post-transcriptionally amplified or downregulated. Classification was based on the difference between protein and RNA logFC values, calculated as delta logFC = protein logFC − RNA logFC, with amplification defined as delta logFC > 0.585 and downregulation defined as delta logFC < −0.585. Categories were further subdivided according to RNA-seq significance (DESeq2, padj < 0.05) and proteomics significance (limma, p-value < 0.05). (F) Gene product counts in the post-transcriptionally amplified and downregulated groups, indicating gene products regulated in males only, females only, or both. (G) KEGG pathway enrichment analysis using ShinyGO v0.77 of gene products post-transcriptionally amplified uniquely in females. The top 10 pathways meeting an FDR < 0.05 threshold are shown and ranked by −log (FDR). (H) Heatmap showing male and female RNA logFC and protein logFC values (HF vs NC) for peroxisomal gene products contributing to the post-transcriptionally amplified Peroxisome KEGG pathway in females. (I) Normalized RNA and protein abundances of selected peroxisomal gene products from (H) in M and F livers under NC and HF diet conditions. Bars represent mean ± s.e.m., with individual mice shown as points. Statistical significance reflects differential expression derived from RNA-seq (DESeq2, padj) and proteomics (limma, p-value) analyses.

Differential expression analysis revealed significant transcriptional remodeling in response to HF diet in both males and females (Figure 3B). Across the male and female datasets, 15,072 total transcripts were identified. Using thresholds of |log FC| > 1 and padj < 0.05, we defined 798 upregulated and 489 downregulated genes in males, and 890 upregulated and 486 downregulated genes in females. Similar to the proteomic results, both sexes exhibited a greater number of upregulated than downregulated genes in response to the HF diet.

Plotting RNA log2 fold changes (RNA logFC) (HF vs NC) in males versus females, we categorized significantly changing genes (|logFC| > 1 and padj < 0.05) into six groups: upregulated in both males and females, upregulated in males only, upregulated in females only, downregulated in both males and females, downregulated in males only, and downregulated in females only (Figure 3C). As observed in the proteomic data, there was partial overlap between sexes (up in both: n = 425; down in both: n = 199); however, a greater number of genes were uniquely regulated within each sex (up only in males: n = 823; up only in females: n = 973; down only in males: n = 566; down only in females: n = 440). These results indicate that HF diet induces both shared and sex-specific transcriptional responses in the liver.

To assess whether transcriptomic changes recapitulate the major metabolic pathway enrichments at the protein level, we performed KEGG pathway analysis across all gene categories defined in figure 3C (Supplementary Figure S3). For gene sets downregulated in both sexes or in males only, pathway enrichments showed partial concordance between transcriptomic and proteomic datasets, with shared representation of metabolic pathways. For example, in both proteomic and RNA-seq data sets, steroid biosynthesis, retinol metabolism, and linoleic acid metabolism were enriched among downregulated pathways in males and females, whereas metabolism of xenobiotics and retinol metabolism were enriched among downregulated male-only pathways. In contrast, for gene sets upregulated in both sexes or uniquely in females, the transcriptomic response was dominated by immune- and inflammation-related pathways rather than the metabolic pathways observed at the protein level. Notably, peroxisomal pathways, which were among the top enrichments in proteins uniquely upregulated in females, were not significantly enriched in the corresponding female-specific transcriptomic category (Figure 3D). These findings indicate that key metabolic adaptations to HF diet in females observed at the proteomic level are not reflected at the transcriptomic level.

Given the significant sex-specific proteome-transcriptome discordances, we next sought to systematically assess the relationship between RNA and protein responses to HF diet. For each gene product detected in both datasets, we plotted the logFC in protein abundance (HF vs NC) against the corresponding RNA logFC within each sex (Figure 3E). Comparison of HF diet–induced RNA and protein logFCs revealed only modest correlations between transcriptomic and proteomic changes in both sexes (males: r = 0.41; females: r = 0.33). To determine whether these correlations exceeded random expectation, we performed permutation testing by randomly scrambling gene-wise RNA–protein pairings within each sex. Scrambled correlations were centered near zero in both males and females, whereas the observed correlations were significantly greater than expected by chance (males: null mean r = −0.0002, empirical p = 1 × 10 ; females: null mean r = 0.0002, empirical p = 1 × 10). Despite being significantly greater than expected by chance, these correlations corresponded to limited explanatory power, with RNA logFC accounting for approximately 17% and 11% of protein logFC variance in males and females, respectively. This lower RNA–protein coupling in females is consistent with the greater discordance between enriched RNA and protein pathways observed prior.

To systematically characterize protein–RNA discordance, we developed a discordance score that enabled classification of gene products as post-transcriptionally amplified or post-transcriptionally downregulated. The score was calculated as the difference between protein and RNA logFCs under HF vs NC conditions (delta logFC = protein logFC − RNA logFC). Gene products were classified as post-transcriptionally amplified when protein abundance increased at least 1.5-fold more than RNA abundance (delta logFC > 0.585), indicating greater protein increase or smaller protein decrease relative to the transcript (RNA did not significantly change and protein increased, RNA increased and protein increased more, RNA decreased and protein did not significantly change, RNA decreased and protein decreased less, and RNA decreased and protein increased). Conversely, gene products were classified as post-transcriptionally downregulated when protein abundance increased at least 1.5-fold less than RNA abundance (delta logFC < −0.585), indicating less protein increase or greater protein decrease relative to transcript (RNA did not significantly change and protein decreased, RNA increased and protein increased less, RNA increased and protein did not significantly change, and RNA increased and protein decreased) (Figure 3E, dashed lines).

Gene product counts in amplified vs downregulated categories revealed that post-transcriptionally regulated responses were largely sex-specific. For post-transcriptional amplification, a greater number of gene products were uniquely amplified in females (n = 282) and males (n = 204), compared to those shared between sexes (n = 56) (Figure 3F). Similarly, for post-transcriptional downregulation, the majority of gene products were unique to each sex (males: n = 118; females: n = 175), with a smaller shared subset (n = 32). These results indicate that post-transcriptional regulation in response to HF diet is predominantly sex-specific, replicating the pattern seen at the protein and RNA levels.

We next performed KEGG pathway enrichment analysis across all categories of post-transcriptionally regulated gene products. Notably, gene products uniquely amplified in females were strongly enriched for peroxisomal pathways (Figure 3G), recapitulating the female-specific enrichment observed at the protein level. Among the remaining categories, only gene products amplified in both sexes and downregulated in males showed significant pathway enrichment (Supplementary Figure S4), with male-specific enrichments partially overlapping with pathways identified at the protein level. To further characterize the female-specific post-transcriptional amplification group, we next examined RNA and protein logFCs (HF vs NC) of specific gene products contributing to the peroxisome enrichment pathway (Figure 3H). The peroxisomal genes uniquely post-transcriptionally upregulated in females belonged to key functional categories of peroxisomal biology, including biogenesis (*Pex3*, *Pex11g*, *Pxmp2*, *Pxmp4*), membrane transporters (*Slc25a17* and *Abcd3*), and β-oxidation (*Ehhadh*, *Hsd17b4*, *Decr2*) and others. Consistent with the classification in figure 3E, females exhibited marked discordance between RNA and protein changes for these gene products, with protein abundance changes exceeding those observed at the transcript level. In contrast, this effect was less pronounced in males.

To visualize the magnitude and significance of RNA and protein changes underlying the logFC analyses, we examined normalized RNA and protein abundances for several representative peroxisomal gene products across all conditions (Figure 3I). In males, these genes exhibited little or no change at either the RNA or protein level under HF diet conditions. In contrast, in females on the HF diet, at the RNA level the same genes showed either no change or a trend toward decreased abundance, whereas their corresponding proteins exhibited statistically significant upregulation. Statistical significance was determined based on RNA-seq (DESeq2, padj < 0.05) and proteomics (limma, p-value < 0.05) HF vs NC comparisons within each sex.

### Direct RNA sequencing identifies sexually dimorphic m6A remodeling in high-fat diet liver

Given the differential metabolic enrichment patterns between transcriptomic analyses and proteomic analyses, we next sought to identify potential mechanisms that might explain this protein–RNA discordance. Protein–RNA discordance can arise through regulation at either the RNA level, such as altered translation efficiency, or at the protein level through changes in protein stability and degradation. To assess whether broad remodeling of protein stability pathways might contribute to the observed amplification phenotype, we examined representative pathways involved in proteasomal degradation, ubiquitination, autophagy, and ER protein processing (Supplementary Figure S5). Neither in males nor in females HF diet appeared to strongly regulate these processes. Therefore, we focused on potential RNA-level regulatory mechanisms.

In recent years, N6-methyladenosine (m6A) has emerged as a widespread regulator of mRNA translation efficiency and post-transcriptional gene expression ^16,32,33^. To understand whether RNA methylation could contribute to discordance between protein and RNA changes in our model system, we investigated m6A regulation on a transcript-by-transcript basis. To characterize m6A dynamics at both single-site and transcript-region resolution, we performed Oxford Nanopore Technologies (ONT) direct RNA sequencing on total RNA extracted from NC and HF diet male and female liver samples (n=3 per condition) (Figure 4A). Principal component analysis (PCA) of filtered m6A counts demonstrated clear separation of samples by sex and diet, with biological replicates clustering tightly within each condition (Figure 4B), indicating high reproducibility of m6A methylation profiles across experimental groups.

**Figure 4:**
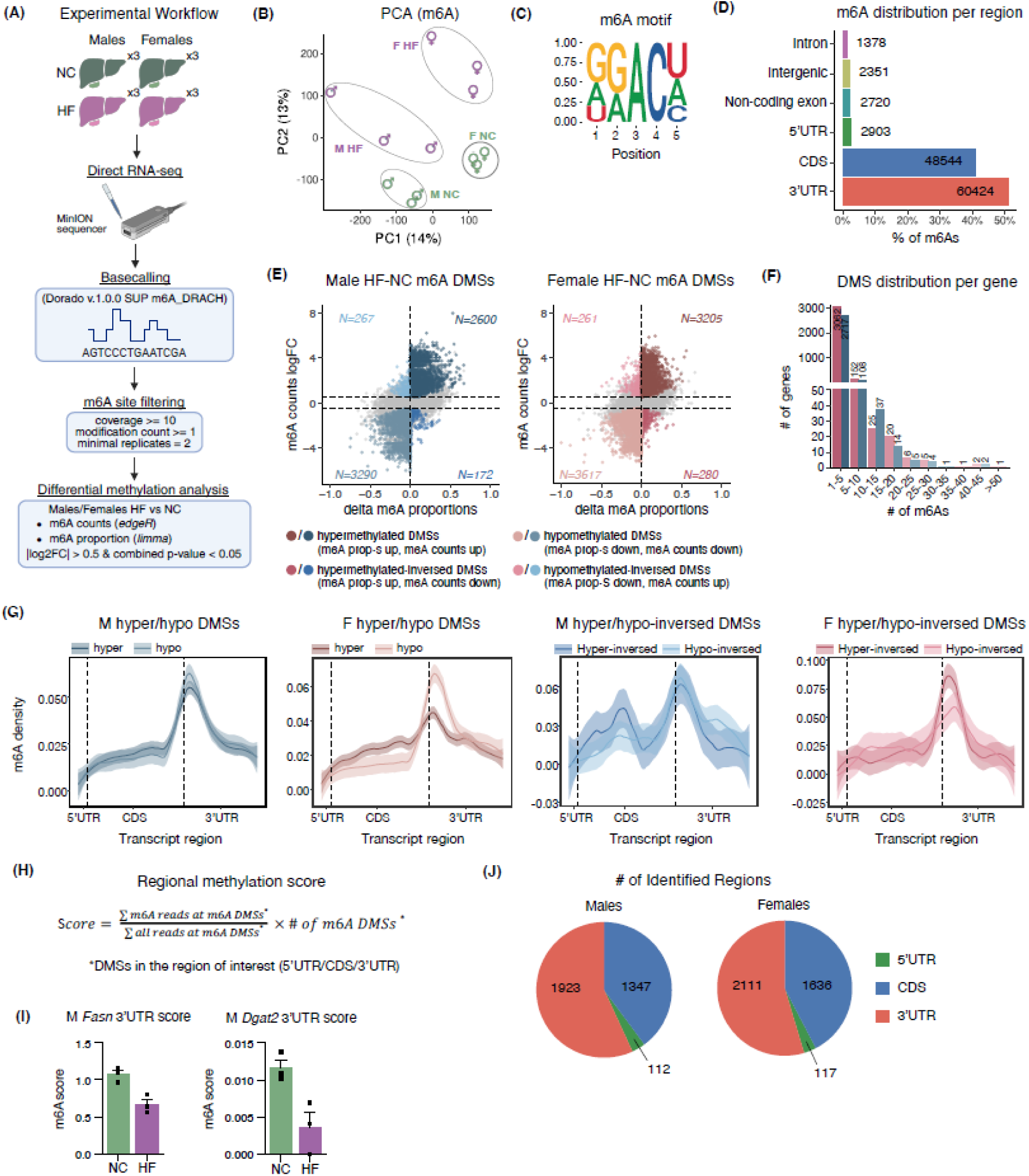
Identification of differentially m6A-methylated sites and regions for male and female livers on HF diet by ONT direct RNA-seq. (A) Schematic overview of m6A site detection and analysis pipeline. (B) Principal component analysis (PCA) of m6A methylation profiles from male (M) and female (F) normal chow (NC) and high fat (HF) diet liver samples (n=3 mice per group). PCAs are based on filtered m6A counts. Each point represents a biological replicate. (C) Motif enrichment around the m6A adenosine (position 3) calculated for filtered m6A sites (m6A DRACH motif). (D) Distribution of filtered m6A sites across transcript regions. (E) Identification and classification of differentially methylated sites (DMSs) between HF and NC conditions in M and F. Scatter plots show changes in m6A proportion (x-axis) versus m6A read count logFC (y-axis) for individual filtered m6A sites. For M HF-NC and F HF-NC comparisons, DMSs classified as hypermethylated (delta m6A proportion > 0, logFC m6A counts > 0.5, combined p-value < 0.05), hypomethylated (delta m6A proportion < 0, logFC m6A counts < -0.5, combined p-value < 0.05), hypermethylated-inversed (delta m6A proportion > 0, logFC m6A counts < -0.5, combined p-value < 0.05), and hypomethylated-inverse (delta m6A proportion < 0, logFC m6A counts > 0.5, combined p-value < 0.05) are colored as indicated. (F) Distribution of numbers of HF-NC m6A DMSs per gene for males and females. (G) m6A meta-profiles showing distribution of HF-NC hyper/hypo-methylated and hyper/hypo-inversed methylated (as defined in figure 4E) HF and NC m6A DMSs across 5’UTR, CDS, and 3’UTR for males (left) and females (right). y-axis represents positional densities (proportions) of m6A DMSs. (H) Formula for calculating a regional (5’UTR/CDS/3’UTR) methylation score. The score represents weighted average of m6A proportions across all DMSs sites in a region of interest (sum of m6A reads divided by sum of all reads at m6A DMSs) multiplied by the number of m6A DMSs in that region. The score is calculated independently for 5’UTR, CDS, and 3’UTR regions of genes in each replicate in each condition. (I) Validation of regional m6A score changes in representative lipogenic genes from the livers of males on HF diet. 3′UTR m6A scores for representative lipogenic transcripts (Fasn and Dgat2) in male liver under NC and HF diet conditions. m6A scores calculated as described in figure 4H. Each point represents an individual biological replicate (n=3). Data are presented as mean ± s.e.m.. (J) Number of 5’UTR, CDS, and 3’UTR regional scores identified for males and females.

To validate the reliability of the detected m6A sites, we first examined sequence motif enrichment surrounding modified adenosines. As expected from the DRACH-aware base calling, the filtered m6A sites were strongly enriched for the canonical m6A DRACH motif (Figure 4C). Predominant localization of m6A sites to the 3’ untranslated region (3’UTR) was also consistent with previously reported transcriptome-wide m6A distributions (Figure 4D).

Having established a high-confidence set of filtered m6A sites, we next identified differentially methylated sites (DMSs) between HF and NC conditions separately for males and females. To increase the rigor, DMSs were classified into four groups based on changes in m6A proportion and m6A read count abundance at the filtered m6A sites between HF and NC conditions: hypermethylated (both proportion of m6A reads and m6A read count increased in HF diet at that m6A site), hypomethylated (both proportion of m6A reads and m6A read count decreased in HF diet at that m6A site), hypermethylated-inversed (proportion of m6A reads increased but m6A read count decreased in HF diet at that m6A site), and hypomethylated-inversed (proportion of m6A reads decreased but m6A read count increased in HF diet at that m6A site) (Figure 4E). Out of 118,320 filtered sites, in both sexes there was a greater number of hypomethylated sites compared to hypermethylated sites (males hypo DMS n=3290 vs males hyper DMS n=2600; females hypo DMS n=3617 versus females hyper DMS n=3205). However, for a smaller subset of sites in which the number of m6A counts and the m6A proportions changed in the opposite direction from each other (hyper-inversed and hypo-inversed DMSs), in females there was a greater number of hypermethylated sites, while for males – hypomethylated (males hypo-inversed DMS n=267 versus males hyper-inversed DMS n=172; females hypo-inversed DMS n=261 versus females hyper-inversed DMS n=280). Such inversed sites could arise when transcript abundance and m6A stoichiometry change independently, causing methylated read abundance and methylation proportion to shift in opposite directions. Across both sexes, the vast majority of genes contained between 1 and 5 m6A DMSs (Figure 4F).

Since m6A modifications located within different transcript regions (5’ untranslated region (5’UTR), coding sequence (CDS), and 3’ untranslated region (3’UTR)) are known to exert distinct regulatory functions, we next examined whether HF diet differentially affects locations of hyper- vs hypomethylated m6A DMS between males and females (Figure 4G). In males, both hyper- and hypomethylated DMSs were predominantly enriched near the beginning of the 3′UTR. In contrast, females exhibited distinct positional patterns depending on methylation directionality. Hypermethylated DMSs in females displayed broader distribution across the CDS and 3′UTR, whereas hypomethylated DMSs showed pronounced enrichment near the beginning of the 3′UTR. Inversed DMS classes exhibited less defined positional distributions in both sexes. This dimorphic positional distribution of hyper versus hypomethylated DMSs further supports distinct effects of HF diet on m6A methylation between males versus females and emphasizes the diet-induced dynamic regulation of m6A in females in particular.

Since our ultimate goal was to determine whether m6A remodeling could contribute to the protein–RNA discordance observed under HF diet conditions, for downstream integrative m6A-RNA-protein analysis we aggregated individual m6A sites into biologically meaningful functional units: 5′UTR, Coding Sequence (CDS), and 3′UTR. The resulting regional methylation score incorporated both changes in average m6A methylation across sites within a region and changes in the number of DMSs present in that region, since both altered methylation stoichiometry and altered site abundance influence the functional impact of m6A remodeling (Figure 4H). As a validation of our approach, we compared regional m6A changes of several key lipogenic genes (*Fasn*, *Dgat2*) to previously reported results of male mice on 4 weeks of HF diet and observed a consistent 3’UTR HF diet-induced demethylation pattern (Figure 4I) ^11^.

Application of this framework identified 112 5′UTR, 1347 CDS, and 1923 3′UTR methylation scores in males, and 117 5′UTR, 1636 CDS, and 2111 3′UTR regional methylation scores in females (Figure 4J). In both sexes, substantially fewer genes contained sufficient 5′UTR DMSs for regional analysis (∼100 genes per sex) compared to CDS and 3′UTR regions. Therefore, subsequent protein–RNA–m6A integration analyses focused primarily on CDS- and 3′UTR-associated methylation dynamics (Figure 5). Together, these results validate identified m6A sites, demonstrate differential regulation of hyper- vs hypomethylated DMSs between males and females on HF, and establish a framework for downstream integrative analysis of m6A changes with protein-RNA discordance. Additionally, although HF diet favored m6A hypomethylation in both sexes, the sex-specific positional distributions of hyper- versus hypomethylated DMSs across transcript regions suggest that male and female livers engage in unique global m6A remodeling programs, with potentially distinct downstream functional outcomes.

**Figure 5.**
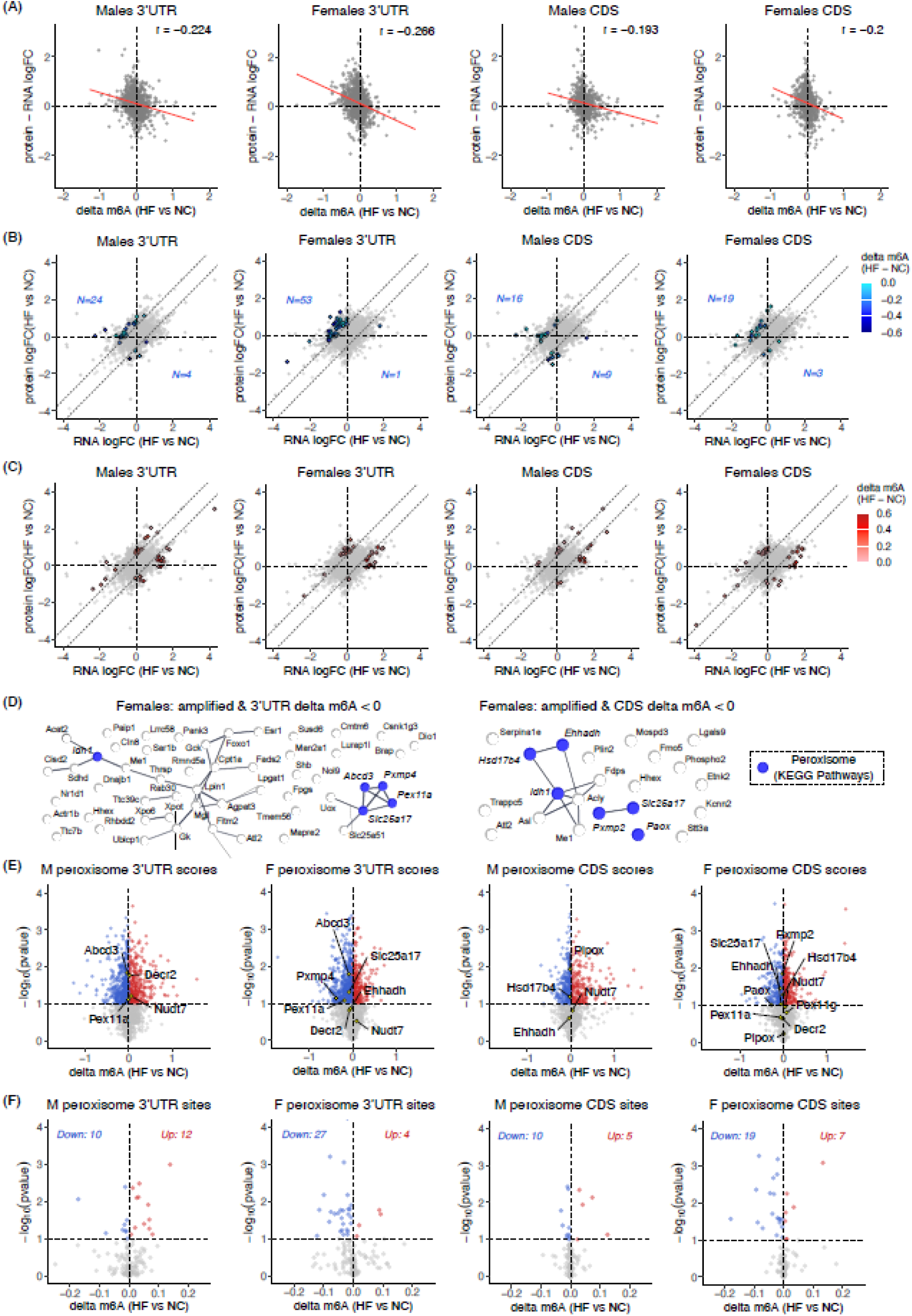
3′UTR m6A hypomethylation of peroxisomal genes in females on HF diet inversely correlates with protein–RNA discordance. (A) Correlation of regional m6A changes, defined as delta m6A = m6A score (HF) − m6A score (NC), in the 3′UTR and coding sequence (CDS) with protein–RNA logFC differences, defined as protein logFC(HF vs NC) − RNA logFC(HF vs NC), in male and female livers. Each point represents a gene detected across the RNA-seq, proteomics, and m6A datasets. Pearson correlation coefficients (r) are indicated. (B) RNA logFC versus protein logFC plots highlighting post-transcriptionally regulated gene products, as defined in figure 3E, that exhibit significantly decreased regional m6A scores on HF diet, defined as delta m6A < 0 and p-value < 0.1, in 3′UTR or CDS regions in males and females. (C) RNA logFC versus protein logFC plots highlighting post-transcriptionally regulated gene products that exhibit significantly increased regional m6A scores on HF diet, defined as delta m6A > 0 and p-value < 0.1, in 3′UTR or CDS regions in males and females. For panels B and C, highlighted points correspond to genes meeting both post-transcriptional regulation and m6A-change criteria, with color gradients reflecting delta m6A values. Dashed lines indicate thresholds used to define post-transcriptional amplification and downregulation. (D) STRING v12.0 network analysis of gene products that are both post-transcriptionally amplified and exhibit decreased m6A score, defined as delta m6A < 0 and p-value < 0.1, in the 3′UTR or CDS, as identified in (B). Genes belonging to the top enriched KEGG pathway, Peroxisome/mmu04146, are highlighted. (E) Volcano plots showing regional delta m6A values in the 3′UTR and CDS for male and female livers under HF versus NC diet conditions. Each point represents a gene, with the x-axis showing delta m6A and the y-axis showing −log(p-value). Significant increases in m6A, defined as delta m6A > 0 and p-value < 0.1, and decreases in m6A, defined as delta m6A < 0 and p-value < 0.1, are indicated in red and blue, respectively. (F) Volcano plots of site-level m6A proportion changes under HF versus NC diet conditions for individual differentially methylated sites within peroxisomal genes of interest: Pex3, Pex11g, Pxmp2, Pxmp4, Slc25a17, Abcd3, Ehhadh, Hsd17b4, Decr2, Nudt7, Pipox,and Paox. Each point represents a single m6A site, with the x-axis showing the change in m6A proportion and the y-axis showing −log(p-value). Significant increases and decreases in m6A proportion, defined as p-value < 0.1, are indicated in red and blue, respectively.

### High-fat diet induces female-specific 3**′**UTR m6A hypomethylation in peroxisomal transcripts associated with protein amplification

To determine whether the previously observed discordances between protein and RNA changes on HF diet could be explained in part by m6A dynamics, we examined the relationship between HF versus NC changes in regional m6A scores (delta m6A) and protein–RNA discordance scores (protein logFC − RNA logFC) across genes. This analysis was performed separately for m6A sites within the 3′UTR and coding sequence (CDS). Across both transcript regions and sexes, delta m6A showed a consistent inverse correlation with protein–RNA discordance (Figure 5A), indicating that decreases in m6A were associated with higher protein logFC relative to RNA. To test whether these inverse relationships exceeded random expectation, we performed permutation testing by randomly scrambling gene-wise pairings between delta m6A and protein–RNA discordance scores within each sex and transcript region. Scrambled correlations were centered near zero in all comparisons, whereas the observed inverse correlations were significantly greater than expected by chance (males 3′UTR: r = −0.224, empirical p = 1 × 10 ; females 3′UTR: r = −0.266, empirical p = 1 × 10 ; males CDS: r = −0.193, empirical p = 1 × 10 ; females CDS: r = −0.200, empirical p = 1 × 10). Notably, correlations were consistently more pronounced in females than in males within both transcript regions, with the strongest inverse relationship observed in the female 3′UTR, aligning with the greater extent of protein–RNA discordance observed in females (Figure 3E). Correlations of delta m6A with RNA logFC and protein logFC independently are depicted in Supplementary Figures S6A and S6B, respectively.

Given that our ultimate goal was to determine whether m6A remodeling could potentially explain the sexually dimorphic protein–RNA discordance, we next examined delta m6A patterns specifically within gene products previously classified as post-transcriptionally amplified or downregulated (Figure 3E). Within these post-transcriptionally regulated gene groups, we highlighted gene products with significantly decreased m6A scores (delta m6A < 0 and p < 0.1; Figure 5B) and significantly increased m6A scores (delta m6A > 0 and p < 0.1; Figure 5C). Among genes with decreased m6A, there was a clear enrichment within the post-transcriptionally amplified category across all conditions. In females, this enrichment was particularly pronounced, with the strongest bias observed in the 3′UTR (amplified: n = 53; downregulated: n = 1), followed by the CDS (amplified: n = 19; downregulated: n = 3). A similar but less pronounced trend was observed in males (3′UTR: amplified n = 24, downregulated n = 4; CDS: amplified n = 16, downregulated n = 9). Notably, the most pronounced enrichment of delta m6A < 0 among post-transcriptionally amplified genes was in the female 3’UTR group, which is consistent with the strongest inverse correlation between protein logFC – RNA logFC vs delta m6A from figure 5A. In contrast, genes with increased m6A did not show a consistent directional bias, with comparable numbers of gene products falling into amplified and downregulated categories across both sexes and regions (Male 3’UTR: 16 amplified, 17 downregulated; Female 3’UTR: 16 amplified, 18 downregulated; Male CDS: 10 amplified, 10 downregulated; Female CDS: 15 amplified, 16 downregulated) (Figure 5C). These results indicate that the relationship between m6A dynamics and post-transcriptional regulation is asymmetric: m6A loss is selectively associated with amplification, while m6A gain does not confer a uniform regulatory outcome.

Since a decrease in m6A score exhibited a clear inverse relationship with discordance of protein vs RNA changes on HF diet, whereas increased m6A did not show a consistent pattern, we next performed pathway enrichment analysis on gene products that were both post-transcriptionally amplified and exhibited a decreased m6A score (Figure 5D). In females, both 3′UTR- and CDS-associated gene sets showed strong enrichment for metabolic pathways, with peroxisomes emerging as the top KEGG pathway in both cases. Several peroxisomal gene products identified in figure 3H were present within these networks, indicating an overlap between m6A remodeling and the female-specific post-transcriptional regulation of protein expression. In contrast, analogous gene sets in males showed limited or no enrichment for metabolically relevant pathways. For males 3′UTR, the top KEGG pathway was “protein processing in endoplasmic reticulum” (mmu04141), while no significant KEGG enrichment was observed for males CDS. Together, these findings indicate that m6A hypomethylation is selectively associated with post-transcriptional amplification of metabolic pathways in females, particularly peroxisomal processes.

Since many gene products in the post-transcriptional amplification group with delta m6A<0 exhibited decreased RNA abundance under HF diet conditions, we performed exon–intron split analysis (EISA) to assess whether negative delta m6A might be associated with RNA stability, instead of our proposed protein–RNA discordance. However, peroxisomal gene products identified in figure 3H and pathway enrichment analyses in figure 5D, demonstrated no significant post-transcriptional RNA stability changes in either sex (Supplementary Figure S7). The only exception was *Nudt7* in females, which exhibited evidence of post-transcriptional downregulation. These results suggest that, for peroxisomal genes, reduced methylation does not affect RNA stability, but more likely post-transcriptionally regulates protein-RNA discordance.

To further examine m6A dynamics within the peroxisomal pathway, we next assessed delta m6A score distributions for all post-transcriptionally amplified peroxisomal gene products identified in figure 3H and figure 5D (except *Idh1*, since it is present not only in peroxisomes, but also in the cytosol). In males, peroxisomal genes did not show a clear directional bias in delta m6A in either the 3′UTR or CDS regions, with both positive and negative values fluctuating around 0 (Figure 5E). In contrast, in females, peroxisomal genes showed a clear bias toward reduced m6A in the 3′UTR, including both significant and non-significant delta m6A score values, indicating a consistent shift toward m6A loss across this pathway. The only exception was *Nudt7*, which exhibited increased m6A. Notably, *Nudt7* was also the only peroxisomal gene showing evidence of post-transcriptional RNA downregulation in EISA analysis (Supplementary Figure S7), suggesting that, in this case, increased m6A may be associated with decreased RNA stability rather than post-transcriptional amplification. In the CDS, females also exhibited enrichment of reduced m6A among significantly changing genes; however, the overall directional bias was less pronounced than in the 3′UTR. Together, these results further support a preferential association between reduced 3′UTR m6A and post-transcriptional amplification of peroxisomal gene products in females.

To resolve m6A dynamics at the single nucleotide level for our genes of interest, we plotted changes of methylated read proportions between HF and NC diets at DMSs across all peroxisomal gene products identified (Figure 5F). In the 3′UTR, females exhibited a pronounced bias toward hypomethylation of DMSs, with the majority of significant sites showing reduced m6A (down: n = 27; up: n = 4), whereas males showed a more balanced distribution between hyper- and hypomethylated DMSs (up: n = 12; down: n = 10). Notably, three of the four hypermethylated sites in females mapped to *Nudt7*, consistent with its behavior being distinct from the rest of peroxisomal gene products in both EISA and region-level m6A analyses. In the CDS, both sexes showed a greater number of hypomethylated than hypermethylated DMSs; however, this bias remained more pronounced in females (down: n = 19 vs males: n = 10). Additionally, the magnitude of m6A reduction at these sites was greater in females compared to males, indicating stronger hypomethylation dynamics under HF diet conditions.

Together with the 3’UTR/CDS methylation score findings, these results demonstrate that peroxisomal genes in females exhibit a coordinated reduction in m6A, particularly within 3′UTR, at both the regional and single-nucleotide levels. This pattern suggests that 3′UTR m6A loss may contribute to enhanced protein output relative to RNA of peroxisomal genes in female livers under HF diet.

## Discussion

Sex differences are increasingly recognized as critical determinants of metabolic disease susceptibility, yet the molecular mechanisms underlying these differences remain incompletely understood. In this study, we validated that female mice are relatively protected from high-fat (HF) diet–induced hepatic steatosis as well as uncovered a distinct adaptive hepatic response at the proteome level, particularly enrichment of peroxisomal pathways. By integrating proteomics, transcriptomics, and Oxford Nanopore direct RNA sequencing, we identify extensive sex-specific epi-transcriptomic remodeling and reveal an association between altered m6A methylation and post-transcriptional amplification of peroxisomal proteins in female liver. Together, these findings support a role for m6A-associated post-transcriptional regulation in sexually dimorphic metabolic adaptation during diet-induced hepatic stress.

Consistent with previous studies on obesogenic diets ^27,34,35^, female mice displayed reduced hepatic lipid accumulation compared to males despite prolonged HF diet exposure. Interestingly, this protection was not readily explained by transcriptional remodeling alone, as female transcriptomic responses were enriched primarily for inflammatory and immune-associated pathways. In contrast, proteomic analyses revealed female-specific metabolic adaptations that may contribute to protection from steatosis, particularly upregulation of proteins involved in peroxisomal biogenesis, transport, and β-oxidation. These changes were not proportionally reflected at the RNA level and were not enriched in males at RNA or protein levels. Together, these findings highlight a sex-specific layer of post-transcriptional regulation that shapes hepatic metabolic adaptation on a high-fat diet.

Peroxisomes play essential roles in hepatic lipid homeostasis; however, the evidence for their pro- versus antisteatotic roles is mixed ^20,25,36,37^. Interestingly, many studies showing that inhibition, not upregulation, of peroxisomal function protects against hepatic steatosis have focused specifically on ACOX1, a peroxisomal enzyme that catalyzes the first step of very long chain fatty acid β-oxidation. However, in our dataset ACOX1 was not significantly upregulated among the high fat diet-induced peroxisomal proteins in females. Thus, the proposed protective role of female peroxisomal remodeling is unlikely to be driven by activation of the same ACOX1-dependent pathways whose inhibition has been previously reported to reduce steatosis. Instead, our findings suggest potential protective roles for alternative peroxisomal pathways, such as ether lipid synthesis or bile acid precursor processing. For instance, plasmalogens, a subclass of ether phospholipids characterized by a vinyl ether bond at the *sn*-1 position, have been strongly implicated in MASLD and hepatic steatosis development in humans and mice alike ^19,24,38,39^. Similarly, the literature reports that increased bile acid synthesis can be protective against hepatic steatosis and that male mice on high fat diet exhibited lowered bile acid levels ^21,40^. Mechanistically, bile acids act as signaling molecules through receptors such as FXR, which modulates hepatic triglyceride metabolism by regulating lipogenesis, lipid oxidation, VLDL handling, and bile acid homeostasis ^41,42^.

A major finding of this study is the identification of extensive sex-specific m6A remodeling in response to HF diet exposure. m6A modifications regulate multiple aspects of RNA metabolism, including translation and transcript turnover, and are increasingly recognized as regulators of nutrient-responsive metabolic remodeling in the liver. Although widespread differential m6A remodeling was observed across transcript regions in both sexes, females demonstrated stronger inverse relationships between m6A changes and protein–RNA discordance scores than males, particularly within the 3′UTR. This finding is consistent with the greater extent of HF diet–induced RNA–protein discordance observed in females relative to males at the global level. Notably, among post-transcriptionally amplified gene products, female livers exhibited coordinated reduction of 3′UTR m6A within peroxisomal transcripts. These female-specific changes were associated with increased protein abundance relative to RNA expression, suggesting that loss of 3′UTR m6A may contribute to enhanced translational output of specific metabolic pathways during HF diet adaptation. Although translation initiation is canonically controlled by 5′UTR features, m6A-dependent translational regulation is not restricted to 5′UTR. Prior work has shown that m6A can promote translation initiation through reader-dependent mechanisms involving 3′UTR m6A sites, while CDS-localized m6As can further influence translation elongation dynamics ^32,43–45^. Thus, the observed female-specific reduction of 3′UTR m6A is consistent with a potential role for regional m6A remodeling in modulating protein output.

Our findings also emphasize the importance of integrating transcriptomic and proteomic datasets when studying hepatic responses to dietary changes, particularly a HF diet. Reliance on transcript abundance alone may overlook biologically important metabolic remodeling that emerges predominantly at the protein level. In this study, transcriptomic responses in females were enriched primarily for inflammatory and immune-associated pathways, whereas proteomic analyses revealed coordinated metabolic adaptations involving peroxisomal pathways. The RNA–protein discordance observed here suggests that post-transcriptional regulation contributes substantially to shaping hepatic metabolic phenotypes during HF diet exposure, with m6A remodeling representing one potential mechanism underlying these adaptive responses. Together, these findings highlight the importance of post-transcriptional remodeling as a component of sex-specific metabolic adaptation.

Several limitations should also be considered. First, although our study identifies strong correlations between m6A remodeling and protein amplification, direct mechanistic experiments are needed to establish causality between specific m6A changes and translational regulation of peroxisomal genes. Second, our analyses were performed using a single HF diet model of MASLD, and future studies across additional dietary and disease models, including Western diet paradigms, will be important for determining the generalizability of these post-transcriptional adaptations. Third, although Oxford Nanopore direct RNA sequencing enabled transcriptome-wide characterization of m6A remodeling at a single nucleotide resolution, limited coverage within 5′UTR regions constrained analyses of m6A changes in transcript regions classically linked to translational regulation.

In summary, our study demonstrates that female protection from HF diet–induced hepatic steatosis is associated with distinct post-transcriptional remodeling, particularly involving peroxisomal pathways and m6A regulation. These findings identify m6A-associated mechanisms as potential contributors to sex-specific metabolic adaptation and highlight the importance of post-transcriptional regulation in shaping hepatic responses to HF diet exposure. More broadly, understanding the regulatory pathways associated with female protection from steatosis may provide insight into general mechanisms governing susceptibility to MASLD and related metabolic disorders.

## ACKNOWLEDGEMENTS

This study was supported in part by grants DK020541 and the 1F30DK145167 from the National Institutes of Health. The Sidoli lab gratefully acknowledges funding from the Hevolution Foundation (AFAR), the ERCM-CFAR Center for AIDS Research, the Aging Biology Foundation, and the NIH Office of the Director (S10OD030286). Research reported in this publication was supported by the Montefiore Einstein Cancer Center Support Grant of the National Institutes of Health under award number P30CA013330. NWS was supported by R35GM156596. KS was supported by a start-up grant from Montefiore Einstein Comprehensive Cancer Center (MECCC) and Albert Einstein College of Medicine, Cancer Center Support Grant 5P30CA013330-53, a Seed grant from the Hirshberg Foundation for Pancreatic Cancer, NIH CTSA Grant 1UM1TR004400 from the Institute of Clinical and Translational Research, Albert Einstein College of Medicine, and a Department of Defense Peer Reviewed Cancer Research Idea Award. RC was supported by T32 AG023475. MH was supported by 5T32GM007491-47. TT and GB received funding from the AIRC Foundation for Cancer Research under MFAG 2020 (ID 24883 project) and AIL Trento.

## Materials and Methods

### Animals and dietary interventions

All animal studies were performed in accordance with protocols approved by the Albert Einstein College of Medicine Institutional Animal Care and Use Committee.

Male and female C57BL/6J mice (Jackson Laboratory, #000664) were obtained at 6 weeks of age and assigned to either a normal chow (NC; LabDiet 5053; 4.5% of calories from fat) or a high-fat (HF; Research Diets, D12492; 60% of calories from fat) diet for 10 weeks. Mice were group-housed in a temperature-controlled room with ad libitum access to food and water under a 12 h light/12 h dark cycle (lights on at 07:00, Zeitgeber time 0).

Body weights were recorded weekly throughout the dietary intervention period. On the day of tissue collection, food was removed at 09:00 and mice were euthanized at 17:00 (8 h fast). Cohorts consisted of n = 4 mice per sex and diet group.

### Tissue collection

For whole liver extraction, mice were anesthetized with isoflurane and euthanized by cervical dislocation. Livers were rapidly excised, cleared of the gallbladder, briefly rinsed in pre-chilled phosphate-buffered saline (PBS), and weighed. Defined portions from consistent anatomical regions of the liver were collected for histological analyses before freezing. The remaining tissue was immediately freeze-clamped using Wollenberger tongs pre-cooled in liquid nitrogen. Frozen liver samples were pulverized under liquid nitrogen using a pre-chilled mortar and pestle and stored at −80 °C until further processing. Aliquots of powdered tissue were used for RNA, protein, triglyceride, metabolomic and m6A analyses.

### Hepatic triglyceride quantification

Hepatic triglyceride content was measured using a colorimetric Triglyceride Quantification Assay Kit (Abcam, ab65336) according to the manufacturer’s instructions with minor modifications. Frozen liver tissue was homogenized as powder under liquid nitrogen, and 25–100 mg of tissue was resuspended in water containing 5% NP-40 substitute (IGEPAL CA-630; Sigma-Aldrich, I3021) at a ratio of 1 mL per 100 mg tissue. Samples were heated at 90 °C for 2–5 min to solubilize lipids and cooled to room temperature twice. Lysates were centrifuged at 12,000 × g for 2 min to remove insoluble material, and the supernatant was collected. Samples were diluted 10-fold in water and further diluted during plate setup to yield a final dilution factor of 50× before assay. Triglycerides were enzymatically hydrolyzed to glycerol using lipase, followed by quantification via a coupled enzymatic reaction generating a colorimetric product. Samples and standards were incubated with reaction mix for 30–60 min at room temperature protected from light, and absorbance was measured at 570 nm using a microplate reader. Triglyceride concentrations were calculated from a glycerol standard curve, adjusted for dilution factors, and normalized to the initial tissue weight. Final values are reported as milligrams of triglyceride per gram of tissue.

### Histology and lipid staining

For histological analyses, defined portions of liver tissue from a consistent anatomical region were collected at the time of dissection. For hematoxylin and eosin (H&E) staining, tissues were briefly rinsed in phosphate-buffered saline (PBS) and fixed in 4% formaldehyde in PBS at room temperature for 24 h in the dark. Following fixation, samples were transferred to 70% ethanol and submitted to the Histology & Comparative Pathology Core for processing, paraffin embedding, sectioning, and staining. Paraffin-embedded liver sections were subjected to standard H&E staining using Harris hematoxylin and eosin following routine deparaffinization, rehydration, and staining procedures.

For lipid staining, fresh liver tissue was snap-frozen in isobutane and processed as frozen sections (8–10 μm). Sections were fixed in buffered formalin, rinsed, and stained with Oil Red O (0.5% in propylene glycol), followed by differentiation in 85% propylene glycol, counterstaining with Mayer’s hematoxylin, and mounting with aqueous medium. Lipid droplets stained red, and nuclei were counterstained blue.

All histological processing and staining were performed by the Histology & Comparative Pathology Core. Research reported in this publication was supported by the Montefiore Einstein Cancer Center Support Grant of the National Institutes of Health under award number P30CA013330.

### Proteomics

#### Sample Preparation and Peptide Loading

Flash-frozen bulk whole liver tissue was used as input material. Tissue was lysed in RIPA buffer supplemented with protease and phosphatase inhibitors and incubated on a rocker for 10 min at 4°C. Lysates were clarified by centrifugation at 14,000 × g for 10 min at 4°C, and protein concentration in the supernatant was measured by BCA assay. Protein input was normalized across samples. Normalized lysates were reduced with dithiothreitol (DTT; final concentration 5 mM) for 1 h at room temperature and alkylated with iodoacetamide (final concentration 20 mM) for 30 min in the dark. Phosphoric acid was then added to a final concentration of 1.2%, and samples were diluted in six volumes of binding buffer (90% methanol, 10 mM ammonium bicarbonate, pH 8.0). Samples were loaded onto S-Trap filters (Protifi), centrifuged at 500 × g for 30 s, and washed twice with binding buffer. Proteins were digested on-column with 1 µg sequencing-grade trypsin (Promega) in 50 mM ammonium bicarbonate at 37°C for 18 h. Peptides were eluted sequentially with (i) 40 µL 50 mM ammonium bicarbonate, (ii) 40 µL 0.1% TFA, and (iii) 40 µL 60% acetonitrile/0.1% TFA. Eluates were pooled, centrifuged at 1,000 × g for 30 s, and dried by vacuum centrifugation. Peptide concentration was measured using the Pierce peptide quantification assay. Immediately prior to LC-MS/MS, peptides were loaded onto Evotips following the Evosep quick-guide workflow: rinse with Solvent B, condition in propanol, equilibrate in Solvent A, load sample, wash with Solvent A, and keep tips wet in Solvent A before acquisition.

#### LC-MS/MS Data Acquisition

Peptides were analyzed using an Evosep LC system (Evosep) coupled to a timsTOF HT mass spectrometer (Bruker Daltonics) with a 20 µm emitter. Peptides were loaded on a PepSep C18 column (15 cm × 150 µm, 1.5 µm) and eluted using Evosep method SPD30. Column temperature was set to 50°C. Data were acquired in dia-PASEF mode with an m/z range of 100-1700 and ion mobility range of 0.6-1.6 1/K0 [V-s/cm²]. Ramp and accumulation times were 100 ms. A total of 32 DIA isolation windows were used across 400-1201 m/z and 0.6-1.6 1/K0 [V-s/cm²]. Collision energy was mobility-ramped from 20 eV at 0.60 1/K0 to 59 eV at 1.6 1/K0.

#### Raw Data Processing

Raw files were searched in DIA-NN (version 2.1.0) using a DIA-NN predicted spectral library generated from the SwissProt mouse protein database (release date: 2024-01-24). Trypsin specificity was used with up to 2 missed cleavages. Carbamidomethylation of cysteine was set as a fixed modification, and methionine oxidation and protein N-terminal acetylation were set as variable modifications. Match-between-runs was enabled, and identification filtering was controlled at 1% FDR.

#### Proteome Informatics and Statistical Analysis

Proteomics data processing and statistical analyses were performed in R (version 4.5.2). The analysis workflow used QFeatures/SummarizedExperiment data structures, followed by feature filtering, transformation, imputation, and differential testing. Proteins were retained when detected in at least three replicates for at least one condition (min_present_per_condition = 3). Intensities were log2-transformed, and missing values were imputed using MinDet (MSnbase::impute, method = "MinDet"). Differential abundance testing was performed using limma with a group-based design matrix and pre-specified contrasts. Results were exported for downstream review and interpretation.

### RNA sequencing

#### RNA extraction and RNA sequencing

Total RNA was isolated from approximately 20 mg of homogenized whole-liver powder using the RNeasy Mini Kit (Qiagen) according to the manufacturer’s instructions. RNA concentration and purity were assessed using a NanoDrop spectrophotometer (Thermo Fisher Scientific). For RNA-sequencing, 1 µg of total RNA per sample was submitted to Azenta Life Sciences for library preparation using the NEBNext Ultra RNA Library Prep Kit (New England Biolabs) followed by sequencing on the NovaSeq platform (Illumina).

#### Raw data processing

Raw sequence data were processed using the nf-core RNA-Seq pipeline (version 3.19.0) ^46^. Reads were aligned to the GRCm39 mouse primary assembly reference genome using the STAR aligner and gene quantification was performed against the GENCODE vM37 primary assembly basic annotation with RSEM to obtain gene expression counts ^47,48^. Expressed genes were determined using zFPKM ^49^. Normalization and differential expression analysis were performed using DESeq2 ^50^.

#### Exon-Intron Split Analysis (EISA)

Exonic and genebody read counts were obtained from STAR-generated BAM files using featureCounts (v2.0.1). Intronic counts were calculated in R (v4.3.3) as the difference between genebody and exonic read counts. Exon–Intron Split Analysis (EISA) was performed using the eisaR package (v1.12.0) with default settings. Analyses were conducted separately for female high-fat diet versus normal chow and male high-fat diet versus normal chow comparisons.

### m6A Profiling (Oxford Nanopore Technologies Direct RNA Sequencing)

#### Library preparation

Total RNA was isolated from homogenized whole-liver powder as described above. RNA concentration and purity were assessed using a NanoDrop spectrophotometer (Thermo Fisher Scientific), followed by evaluation of RNA integrity using the Agilent 2100 Bioanalyzer (Agilent Technologies). Only samples with an RNA integrity number (RIN) greater than 7 were used for downstream analysis. For Oxford Nanopore Technologies (ONT) direct RNA sequencing, ∼3500 ng of total RNA per sample was used as input for library preparation using the Direct RNA Sequencing Kit (SQK-RNA004; ONT) according to the manufacturer’s instructions. Libraries were sequenced individually on FLO-MIN004RA flow cells using the MinION MK1D platform (ONT). Raw nanopore signal data were used for downstream m6A detection and differential methylation analyses.

#### Data Processing

Raw pod5 files were basecalled using Dorado (v1.1.1) with models rna004_130bps_sup@v5.2.0 and rna004_130bps_sup@v5.2.0_m6A_DRACH@v1. The - modified-bases-threshold was left at the standard 0.05. Reads were aligned to GRCm39 using Dorado aligner (v.1.1.1) and minimap2, specifying splice-aware mode. m6A modifications were extracted from the aligned bam files using modkit (V0.5.0) specifying base modification pass threshold values at 0.9. Modification counts and proportion tables were subsequently analyzed with R (v.4.4.2). Sites with minimum coverage of 10 reads and at least one modified call in two out of three replicates of a condition were retained. Differentially methylated m6A sites were identified combining differential testing on modification counts and frequencies. Negative binomial regression was employed on modification counts, using the glmQLFit function from EdgeR, while linear regression with empirical Bayes statistics was employed on modification proportions using limma lmFit and eBayes functions. Changes between High-Fat Diet and Normal-Chow in Males and Females were tested (Males HF vs NC and Females HF vs NC) in both cases. To identify high-confidence differences and define condition-associated classes to each site, we computed a combined p-value using Fisher’s method. Modification sites with m6A counts |log2FC| ≥ 0.5, a change in m6A proportions in the same direction, and a combined p-value ≤ 0.05 were considered as consistently hypermethylated or hypomethylated. Modification sites with m6A counts |log2FC| ≥ 0.5, a change in m6A proportions in the opposite direction, and a combined p-value ≤ 0.05 were considered as hypermethylated-inversed or hypomethylated-inversed. Sites aggregation at the transcript level was performed by applying the following formula:

m6A modification score = (∑GeneN m6A counts)/(∑GeneN m6A coverage) × number of called m6As GeneN. To compute gene m6A modification score, we considered only the differentially methylated sites previously identified.

### Multi-omics Integration

RNA-seq, proteomics and m6A datasets were integrated at the gene level using gene symbols. For RNA–protein analyses, genes were retained if they were detected in both datasets and had non-missing log2 fold-change and statistical values. For RNA–protein–m6A integration, genes were retained only if they were present in the RNA-seq, proteomics and corresponding regional m6A score datasets. Analyses were performed separately by sex and, for m6A integration, separately for 5′UTR, CDS and 3′UTR regions. Pearson correlations were calculated in R using cor.test. RNA–protein correlations were calculated separately by sex. Correlations between delta m6A and protein–RNA discordance were calculated separately by sex and transcript region. Genes with altered regional m6A were defined using delta m6A < 0 or >0 with nominal P < 0.1 by t-test and were intersected with post-transcriptionally amplified or downregulated gene sets for downstream pathway analysis.

### Functional enrichment and network analysis

KEGG pathway enrichment was performed using ShinyGO v0.77 with mouse selected as the species. For RNA-seq-derived lists, genes detected in the RNA-seq dataset were used as background. For proteomics-derived lists, proteins detected in the proteomics dataset were used as background. Enrichment was performed using an FDR cutoff of 0.05 and pathway size range of 2–2,000 genes.

Protein network analyses were performed using STRING v12.0 with *Mus musculus* as the organism. Networks were generated using the full STRING network with all interaction sources enabled and a medium confidence interaction score cutoff of 0.400. STRING enrichment terms were displayed using FDR ≤ 0.05 and a minimum count of two proteins per term. KEGG pathway annotations from STRING were used to highlight proteins belonging to selected enriched pathways.

### Data visualization and statistical analyses

Data processing, statistical analyses and visualization were performed using R, GraphPad Prism, Microsoft Excel and Fiji/ImageJ. Figures were assembled using BioRender, Inkscape and Microsoft PowerPoint.

## Code and data availability

The code for data processing is available in Github at https://github.com/maxhorton13/multi-omic-analysis-of-sex-differences-in-hepatic-steatosis. The RNA-seq data are available in the GEO repository at GSE335522. The proteomics raw files and result files are available on ProteomeXchange (PRIDE) under the project number PXD079603. For reviewers, please access the files using the token: HGKTrRkg1eFI.

## Supplementary Material

**Supplementary Figure S1.**
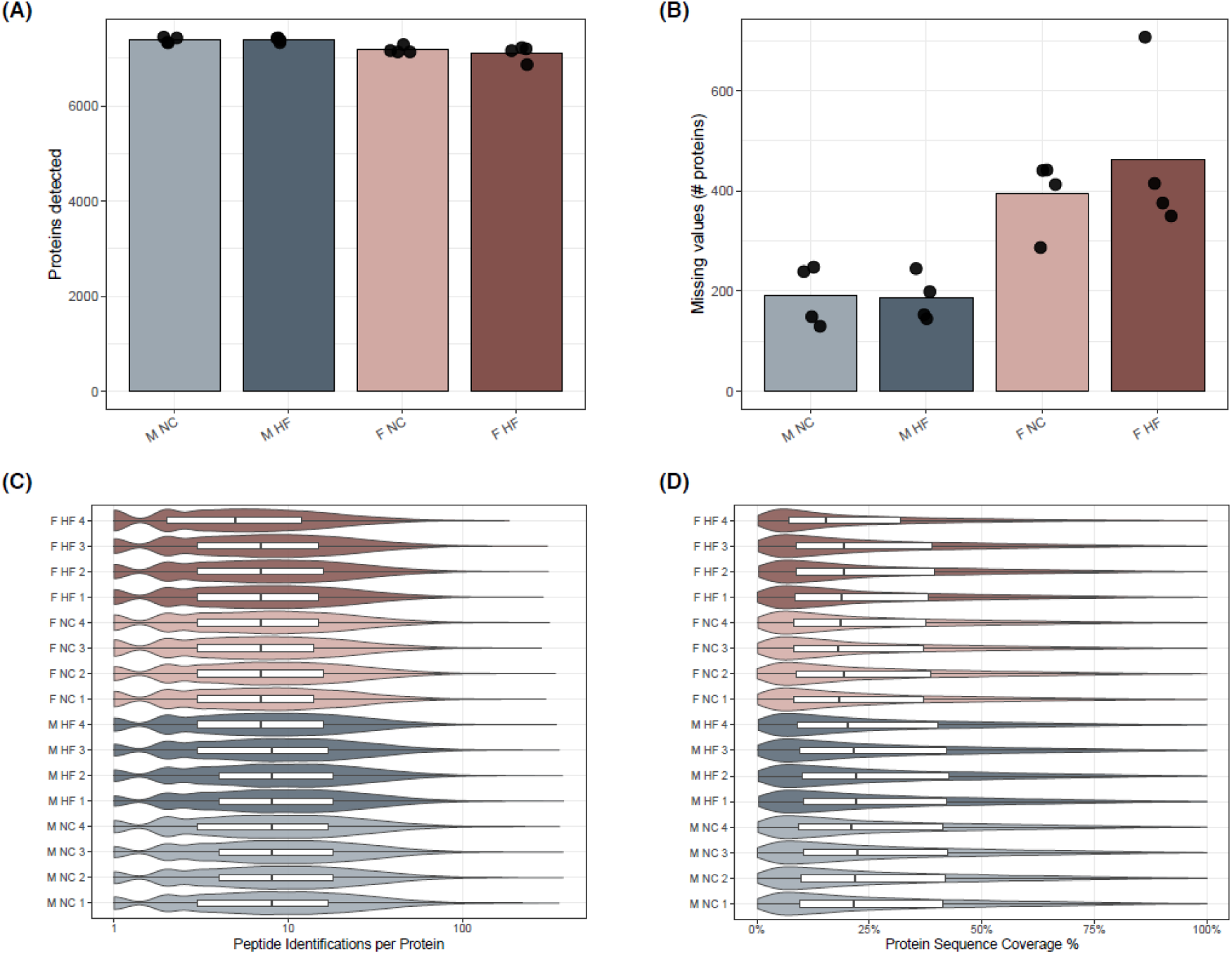
Proteomics preprocessing and quality control. (A) Number of proteins detected in each sex and diet condition. Bars represent the mean across biological replicates, with individual samples shown as points. (B) Number of missing protein abundance values per condition. Bars represent the mean across biological replicates, with individual samples shown as points. (C) Distribution of peptide identifications per protein across individual samples. (D) Distribution of protein sequence coverage percentages across individual samples. For panels C and D, violin plots show sample-level distributions with embedded boxplots indicating the median and interquartile range.

**Supplementary Figure S2:**
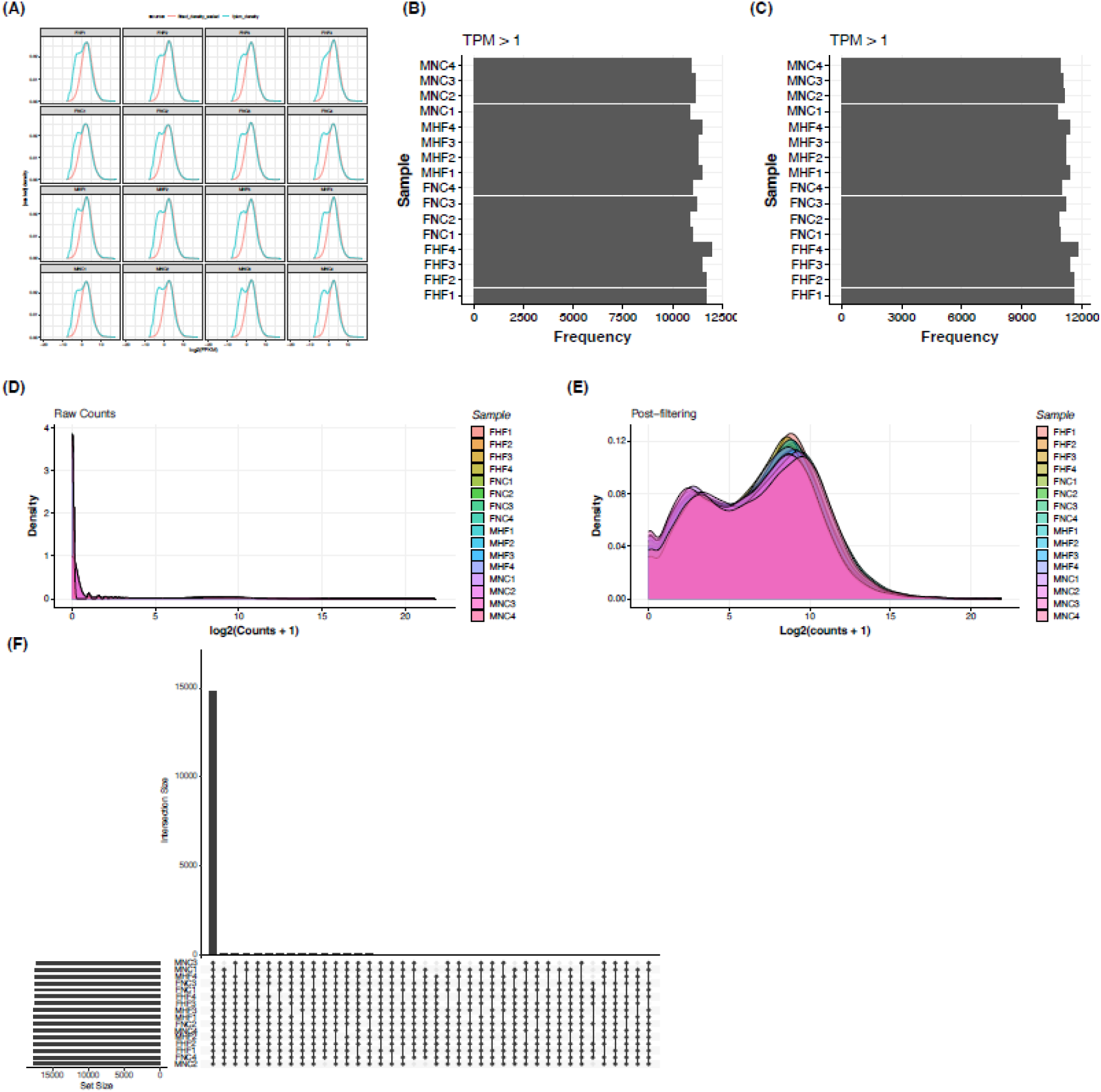
RNA-sequencing preprocessing and quality control. (A) zFPKM-based expression filtering across RNA-seq samples. Blue curves indicate empirical zFPKM distributions, and red curves indicate fitted reference distributions used to identify expressed genes for downstream analyses. (B) Number of genes detected per sample before filtering using a TPM > 1 threshold. (C) Number of genes retained per sample after expression filtering using a TPM > 1 threshold. (D) Distribution of raw gene-level RNA-seq counts across all samples prior to filtering. (E) Distribution of gene-level RNA-seq counts across all samples after filtering. For panels D and E, counts are shown as log (counts + 1). (F) UpSet plot showing the overlap of expressed genes across RNA-seq samples after filtering. Bar heights represent intersection sizes, and connected points indicate the samples included in each intersection.

**Supplementary Figure S3:**
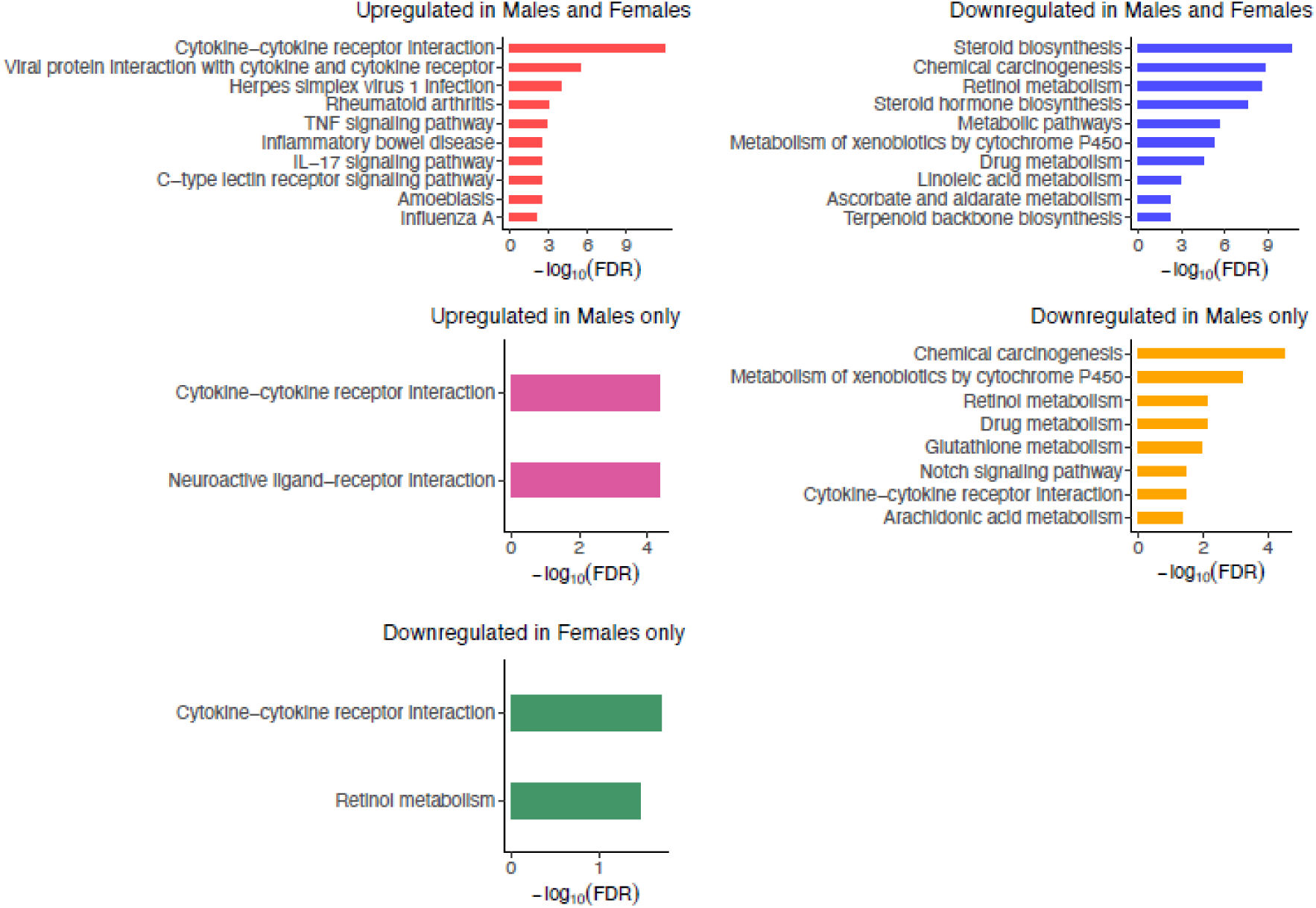
Transcriptomic pathway enrichments across sex-specific and shared gene sets. KEGG pathway enrichment analysis (ShinyGO v0.77) of differentially expressed genes identified in figure 3C, stratified into categories based on sex-specific and shared regulation (upregulated in males and females, downregulated in males and females, upregulated in males only, downregulated in males only, and downregulated in females only). Top enriched pathways meeting an enrichment threshold of FDR < 0.05 are shown (maximum of 10 per category) and ranked by –log(FDR). The category “upregulated in females only” is presented in Figure 3D.

**Supplementary Figure S4:**
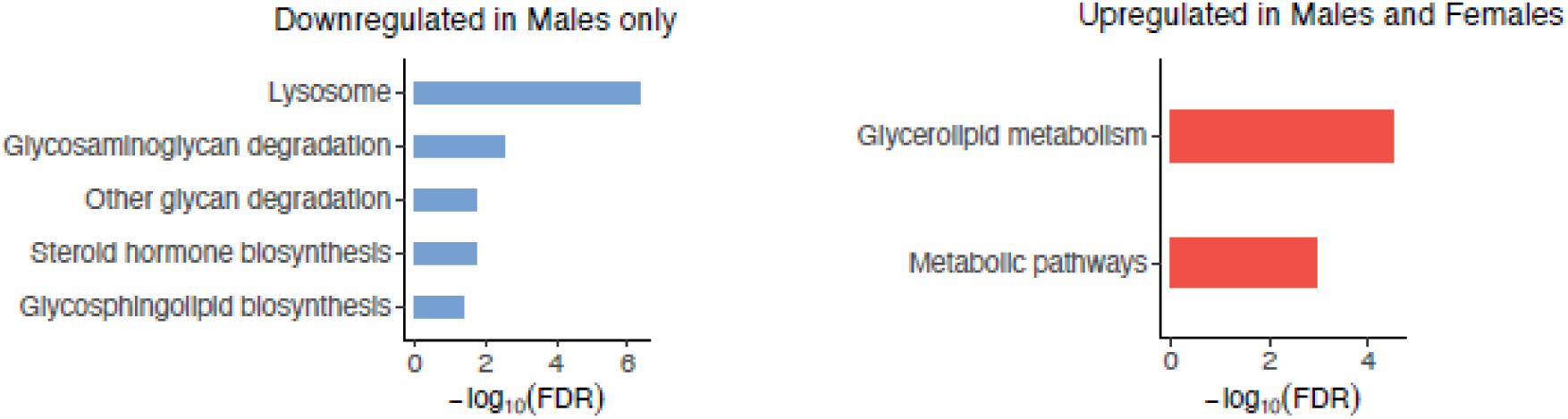
Pathway enrichment of post-transcriptionally regulated gene sets. KEGG pathway enrichment analysis (ShinyGO v0.77) of gene sets classified based on post-transcriptional regulation (Figures 3E–F). Shown are categories exhibiting significant enrichment (FDR < 0.05), including gene products post-transcriptionally downregulated uniquely in males (left) and post-transcriptionally amplified in both male and females (right). Pathways are ranked by –log(FDR).

**Supplementary Figure S5:**
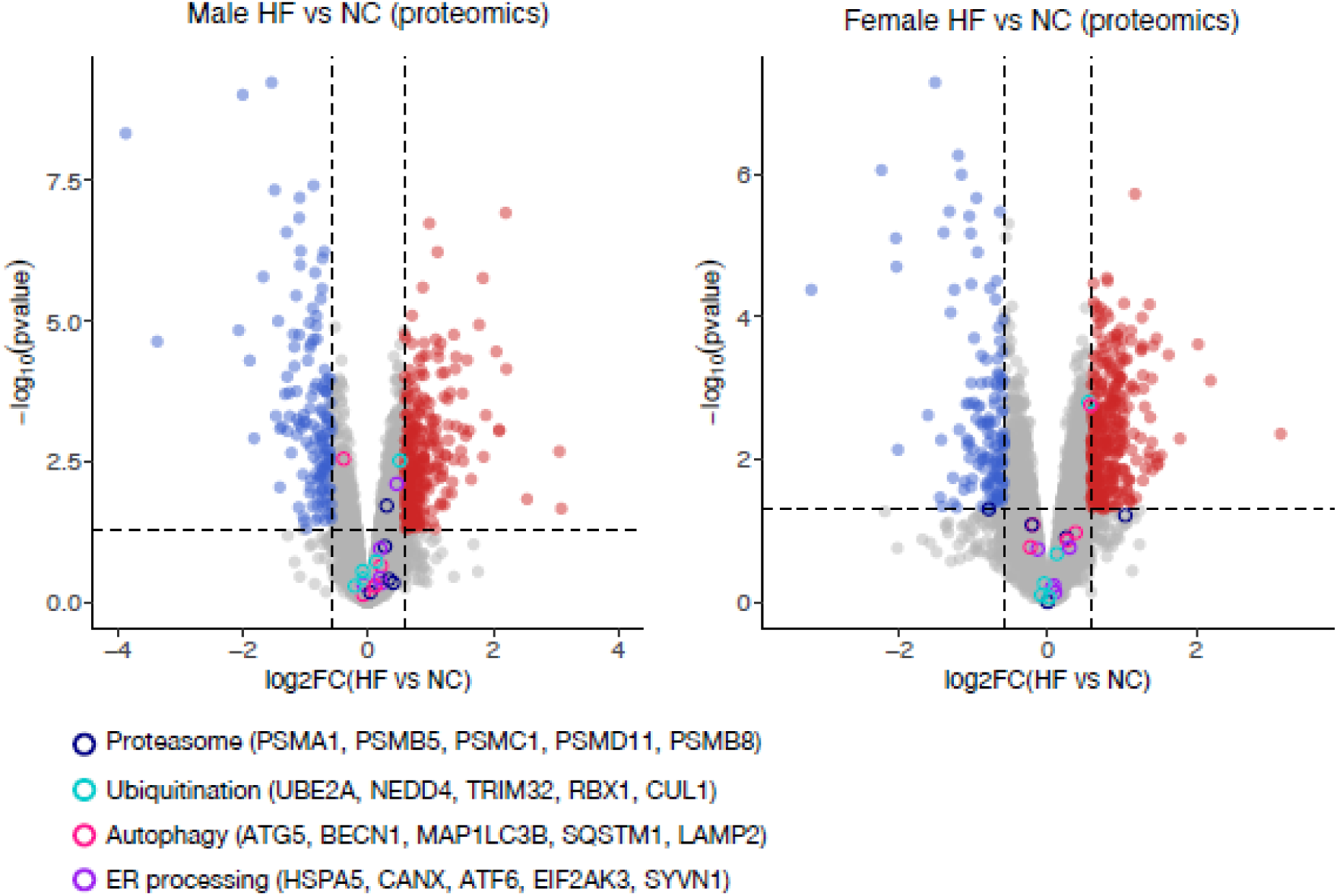
Proteostasis pathways in the liver from mice fed a HF or NC diet. Volcano plots showing differential protein abundance in males and females on HF vs NC diets. Differential expression was calculated using limma. Significance threshhold is |logFC| > 0.585 and p-value < 0.05. Representative proteins involved in protein stability pathways (proteasomal degradation, ubiquitination, autophagy, and ER protein processing pathways) are colored according to the legend.

**Supplementary Figure S6.**
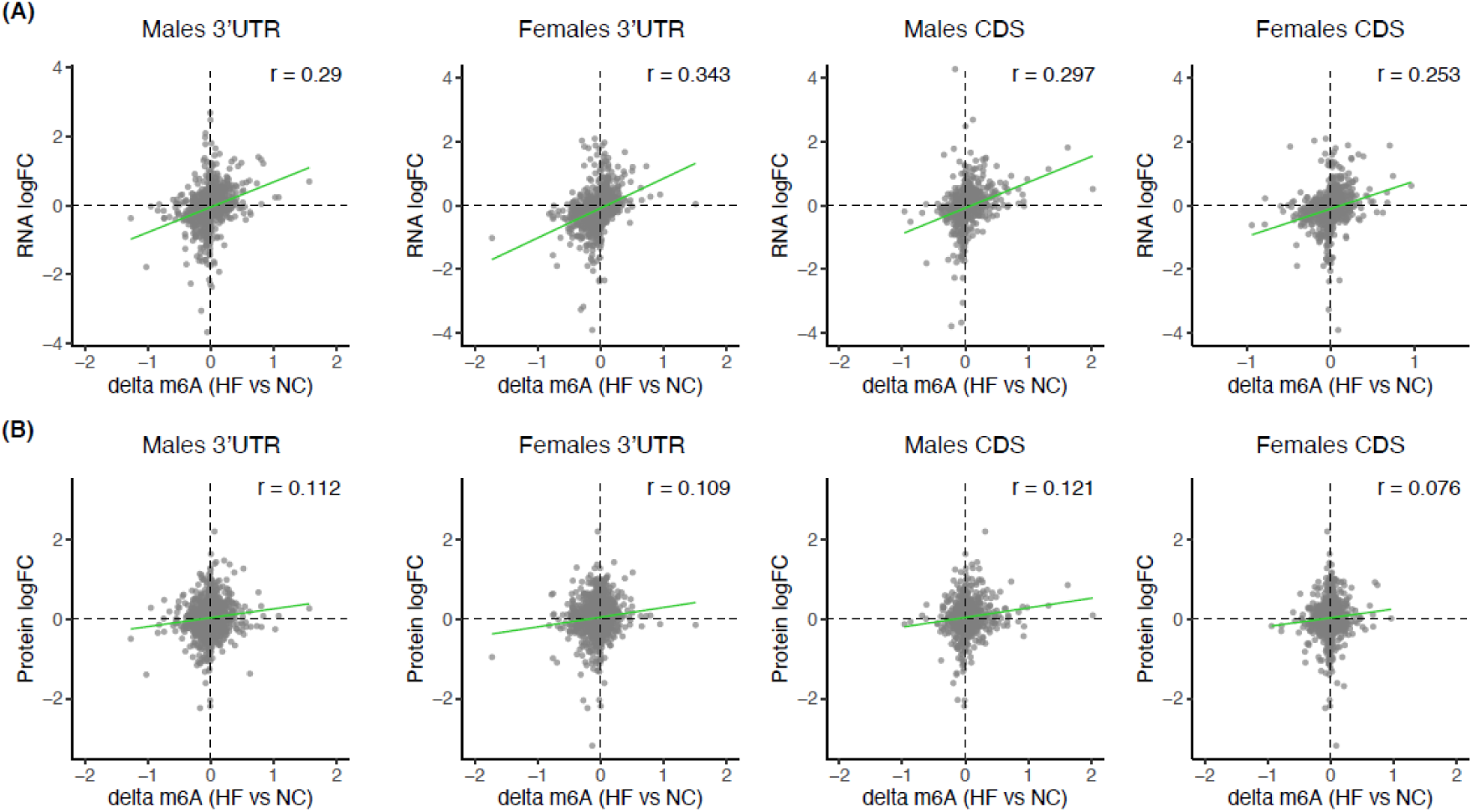
Correlations of delta m6A with RNA and protein logFCs across transcript regions. (A) Correlation of regional m6A score changes, defined as delta m6A = m6A score(HF) − m6A score(NC), with RNA logFC in male and female livers across 3′UTR and coding sequence (CDS) regions. (B) Correlation of regional m6A score changes with protein logFC across the same sex- and transcript-region-specific comparisons. Each point represents a gene detected across RNA-seq, proteomics, and m6A datasets, using the same gene sets as in figure 5A. Pearson correlation coefficients (r) are indicated.

**Supplementary Figure S7:**
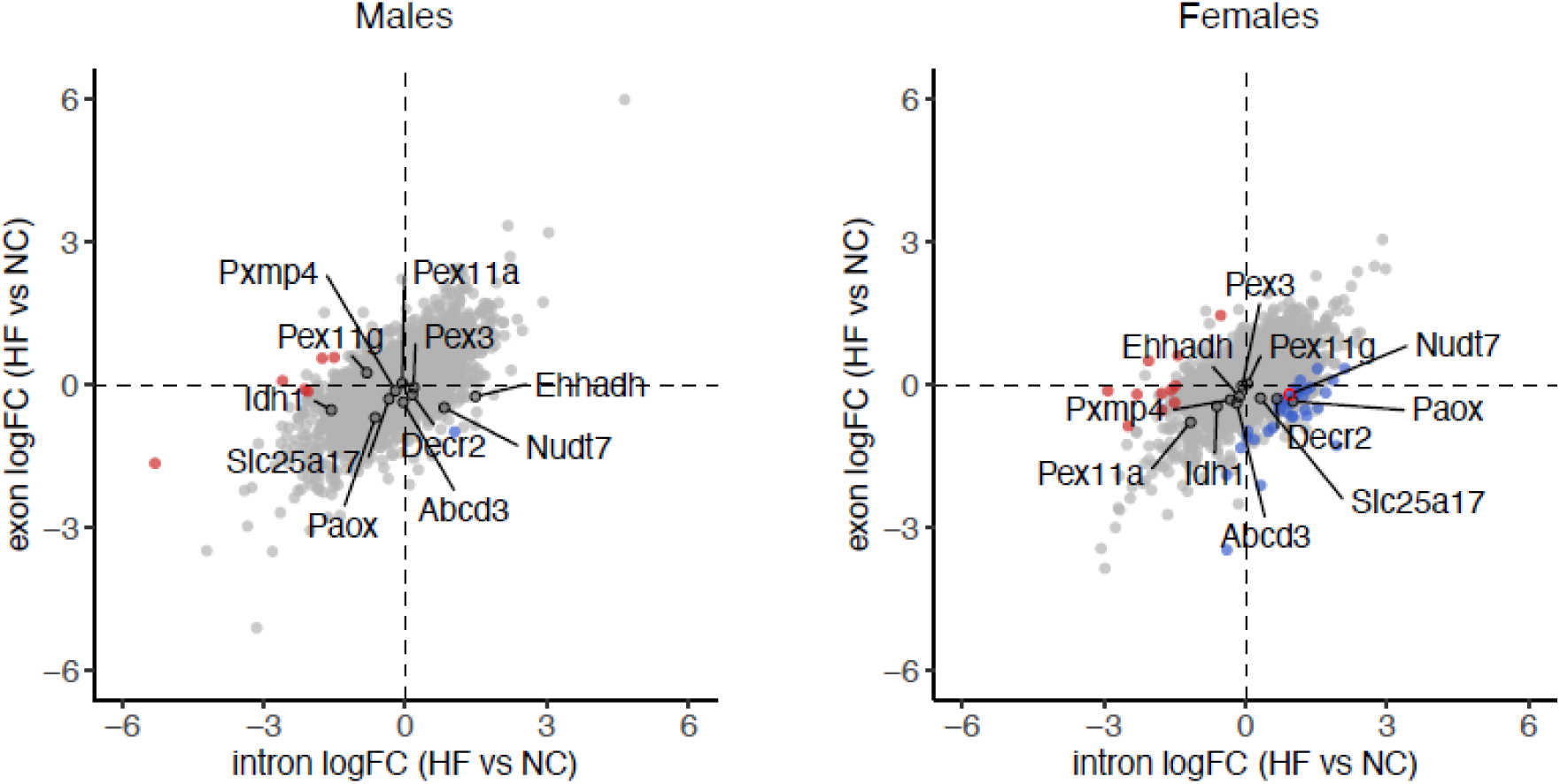
Exon–intron split analysis (EISA) for post-transcriptionally amplified peroxisomal genes. EISA comparing exon and intron log2 fold changes (HF vs NC) in male and female livers. Exonic and intronic read counts were derived from RNA-seq data and quantified separately as previously described ^51^. Each point represents a gene, with the x-axis indicating intron logFC (proxy for transcriptional regulation) and the y-axis indicating exon logFC (combined transcriptional and post-transcriptional effects). Genes deviating from the diagonal reflect post-transcriptional regulation. Genes exhibiting significant post-transcriptional regulation (|exon − intron logFC| > 1, FDR < 0.05) are highlighted (red, post-transcriptionally upregulated; blue, post-transcriptionally downregulated). Peroxisomal genes of interest (as defined in figure 3H and figure 5D) are labelled.

## Notes

### Competing Interest Statement

The authors have declared no competing interest.

